# The novel lncRNA *lnc-NR2F1* is pro-neurogenic and mutated in human neurodevelopmental disorders

**DOI:** 10.1101/410837

**Authors:** Cheen Euong Ang, Qing Ma, Orly L. Wapinski, Shenghua Fan, Ryan A. Flynn, Bradley Coe, Masahiro Onoguchi, Victor H. Olmos, Brian T. Do, Lynn Dukes-Rimsky, Jin Xu, Qian Yi Lee, Koji Tanabe, Liangjiang Wang, Ulrich Elling, Josef Penninger, Kun Qu, Evan E. Eichler, Anand Srivastava, Marius Wernig, Howard Y. Chang

**Affiliations:** Institute for Stem Cell Biology and Regenerative Medicine and Department of Pathology, Stanford University, Stanford, CA 94305, USA; Department of Bioengineering, Stanford University, Stanford, CA 94305, USA; Center for Personal Dynamic Regulomes, Stanford University, Stanford, CA 94305, USA; Department of Dermatology and Department of Genetics, Stanford University, Stanford, CA 94305, USA; J.C. Self Research Institute of Human Genetics, Greenwood Genetic Center, Greenwood, S.C. 29646; Howard Hughes Medical Institute and Department of Genome Sciences, University of Washington, Seattle, WA 98195; Department of Genetics and Biochemistry, Clemson University, Clemson, SC 29634; Institute of Molecular Biotechnology of the Austrian Academy of Science (IMBA), Vienna Biocenter (VBC), Dr. Bohr Gasse 3, 1030 Vienna, Austria; Present address: CAS Key Laboratory of Innate Immunity and Chronic Diseases, School of Life Sciences and Medical Center, University of Science and Technology of China, Hefei 230027, China

## Abstract

Long noncoding RNAs (lncRNAs) have been shown to act as important cell biological regulators including cell fate decisions but are often ignored in human genetics. Combining differential lncRNA expression during neuronal lineage induction with copy number variation morbidity maps of a cohort of children with autism spectrum disorder/intellectual disability versus healthy controls revealed focal genomic mutations affecting several lncRNA candidate loci. Here we find that a t(5:12) chromosomal translocation in a family manifesting neurodevelopmental symptoms disrupts specifically *lnc-NR2F1*. We further show that *lnc-NR2F1* is an evolutionarily conserved lncRNA functionally enhances induced neuronal cell maturation and directly occupies and regulates transcription of neuronal genes including autism-associated genes. Thus, integrating human genetics and functional testing in neuronal lineage induction is a promising approach for discovering candidate lncRNAs involved in neurodevelopmental diseases.

---

Eukaryotic genomes are extensively transcribed to produce long non-coding RNAs (lncRNAs) in a temporally and spatially regulated manner^1^. Until recently, lncRNAs were often dismissed as lacking functional relevance. However, lncRNAs are emerging as critical regulators of diverse biological processes and have been increasingly associated with a wide range of diseases, based primarily on dysregulated expression^2^. LncRNAs represent a new layer of complexity in the molecular architecture of the genome, and strategies to validate disease relevant lncRNAs are much needed. High-throughput analyses have shown that lncRNAs are widely expressed in the brain and may contribute to complex neurodevelopmental processes^2-9^. However, few studies have examined the role of lncRNAs in brain development mostly due to technical difficulties. Direct lineage conversion by the transcription factors <u>B</u>rn2, <u>A</u>scl1 and <u>M</u>yt1l (termed BAM factors in combination) into induced neuronal (iN) cells, recapitulates significant events controlling neurogenesis programs^10-12^, and therefore, it is a facile and informative system to study the role of lncRNAs in the establishment of neuronal identity.

The noncoding genome has emerged as a major source for human diversity and disease origins. Given that less than 2% of the genome encodes protein-coding genes, the majority of the genomic landscape is largely encompassed by non-coding elements. Efforts to identify genetic variation linked to human disease through genome-wide association studies revealed a significant majority affecting the non-coding landscape. Based on their expression and diversity in the mammalian brain, we postulate neuronal lncRNAs may be recurrently affected by mutations that disrupt normal brain function. Neurodevelopmental disorders manifest as a spectrum of phenotypes particularly early in life ^13^. Recent studies suggest that this diversity is the result of different combinations of mutations in multiple genes, often impacting key pathways such as synapse function and chromatin regulation. Nonetheless, despite recent findings that have greatly increased the number of protein coding genes implicated in human intellectual disability and autism, a majority of patients lack well-understood genetic lesions which include a large number of inherited variants occur in noncoding regions that could not be interpreted ^14-21^.

In this study, we used an integrative approach to identify lncRNA genes important for human disease by incorporating high throughput cell fate reprogramming, human genetics, and lncRNA functional analysis. In addition, we developed a pipeline to enrich for lncRNAs with neuronal function and are associated with disease through focal mutations in patients with autism spectrum disorder and intellectual disability (ASD/ID). Furthermore, we show that one of these lncRNAs, *lnc-NR2F1* participates in neuronal maturation programs *in vitro* by regulating the expression of a network of genes previously linked to human autism.

## Results

### LncRNA candidate loci are recurrently mutated in patients with neurodevelopmental disorders

LncRNAs have been associated with human diseases primarily through alterations in expression levels ^22-24^. However, little is known about mutations affecting the genomic loci that encode lncRNAs. We used RNA sequencing data from profiling transcriptomic changes during induced neuronal cell reprogramming at various time points to enrich for lncRNA genes potentially involved in neural fate specification. Of the 27,793 known and predicted lncRNAs, 287 were differentially expressed among any two time points (0, 2, 13 and 22 days) in our dataset (>2-fold, p<0.05). After imposing multiple criteria based on their expression during iN reprogramming and brain development, chromatin association and their proximity to neuronal genes, we shortlisted 35 candidate lncRNAs further analyses. (**Supplementary text 1, Fig. S1A-H, S2**).

We next interrogated these 35 mouse lncRNA loci in patients with autism spectrum disorder and intellectual disability (ASD/ID). Firstly, we found that 28 of the 35 mouse lncRNA candidates have human synteny, and 10 loci were already annotated as non-coding RNAs (**Fig. 1A and S3A**). We next overlapped the 28 human lncRNA candidates and the remaining iN cell-lncRNAs coordinates to a CNV morbidity map recently built from 29,085 patients diagnosed with a spectrum of neurodevelopmental disorders and craniofacial congenital malformations, and 19,584 controls ^25,26^. This approach was motivated by the fact that the CNV morbidity map has successfully identified novel syndromes characterized by recurrent mutations affecting protein-coding genes of ASD/ID patients, and has offered mechanistic insight into the drivers of the pathogenesis ^25-27^.

**Fig. 1.**
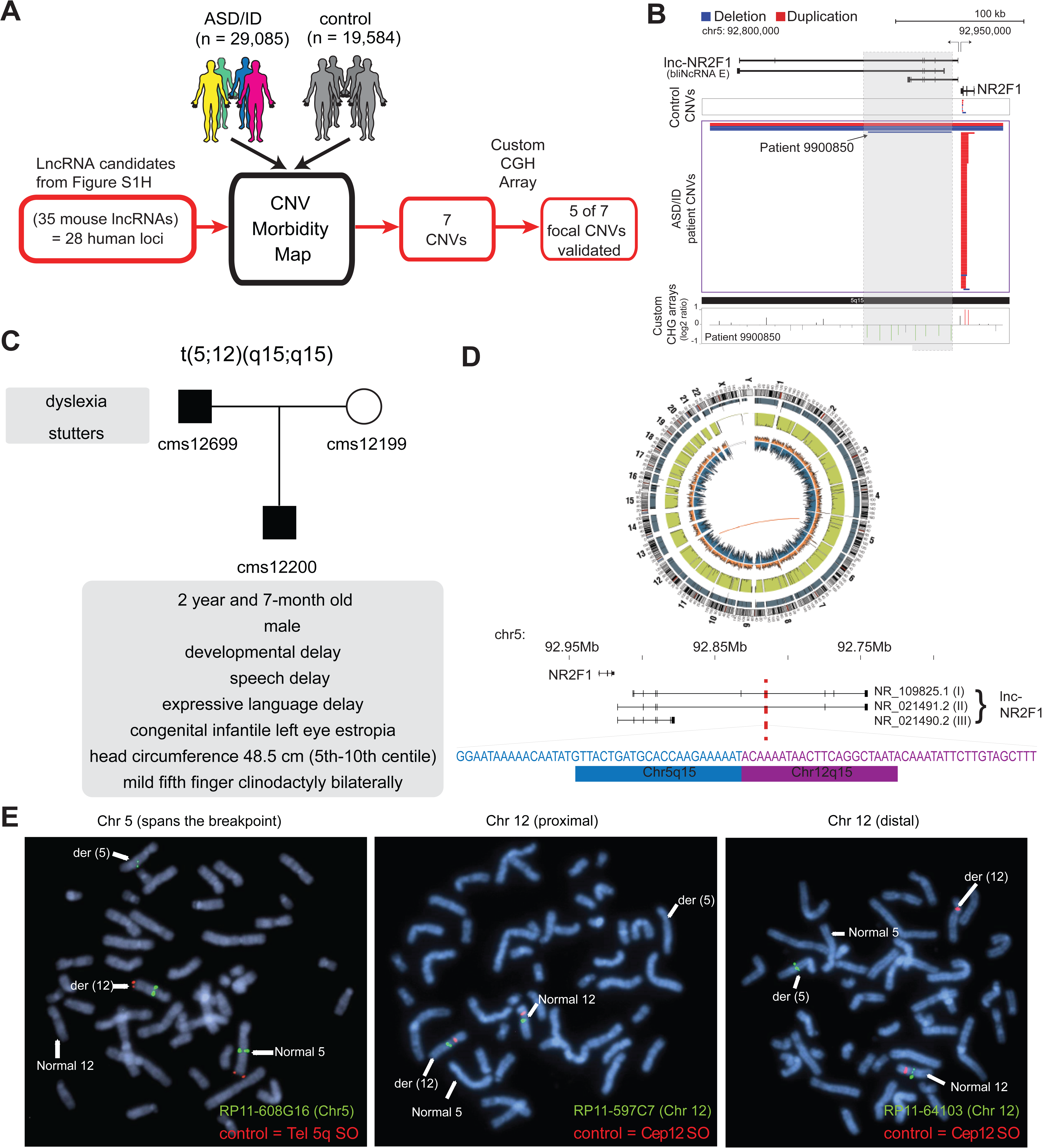
lncRNA loci are recurrently mutated in patients with neurodevelopmental disorders. (A) Schematic representation of CNV morbidity map analysis for candidate lncRNAs and all other iN lncRNAs loci. The 35 mouse lncRNA candidates (28 human loci) is from Figure S1H. (B) Top: Representative tracks for lncRNA E locus, also known as *lnc-NR2F1*. Depicted in blue are deletions and in red duplications. Arrow points to patient with focal deletion affecting the *lnc-NR2F1* locus only. Bottom: Custom CGH arrays used to validate chromosomal aberration in patient 9900850 harboring focal deletion represented in green signal. (C) Genetic pedigree analysis for family with paternally inherited balanced chromosomal translocation (5;12)(q15;q15), including a summary of clinical features for patient CMS12200 and father. The mother has a normal karyotype. Listed in the box are the symptoms of the patients. (D) Top: Circa plot representing the pathogenic chromosomal event for patient CMS12200 involving chromosomes 5 and 12. Bottom: Representative chromosome ideogram and track of the balanced chromosomal break affecting patient CMS12200. Below the ideoplot is the schematic representation of predominant human isoforms for *lnc-NR2F1* and the site of the break site disrupting the long isoforms. (E) The locations of the probes are in supplementary figure S4C. Left: Metaphase spread from patient CMS12200 with the t(5;12) translocation showing FISH signals obtained with the clone RP11-608G16 (green) spanning Chromosome 5 breakpoint, and a Chromosome 5 telomere-specific probe (red). Middle: Metaphase spread from patient CMS12200 with the t(5;12) translocation showing FISH signals obtained with the clone RP11-597C7(green) proximal to Chromosome 12 breakpoint, and a Chromosome 12 centromere-specific probe (red). Right: Metaphase spread from patient CMS12200 with the t(5;12) translocation showing FISH signals obtained with the clone RP11-641O3 (green) distal to Chromosome 12 breakpoint, and a Chromosome 12 centromere-specific probe (red).

Intersecting genomic coordinates of human lncRNA candidate loci to the CNV morbidity map revealed 7 focal CNVs enriched in disease that overlap with 5 candidate lncRNA loci: E (FLJ42709), H (LOC339529), Z (LOC100630918), D (LINC00094) and O (LO467979) (Sequences in supplementary documents). Among these 7 focal CNVs, 5 events corresponded to small deletions in human lncRNA candidates E, H, Z, D and O. Two lncRNA loci corresponding to lncRNAs H and Z were affected by two independent and different focal CNVs. We verified that all five human lncRNAs are expressed during human brain development (**Fig. S3A**). We then designed a custom tilling array with dense coverage of the affected loci for comparative genomic hybridization (CGH) to validate the focal CNVs in the genomic DNA from affected individuals. 5 of 7 focal CNVs affecting the lncRNA loci were tested and validated (**Fig. 1A, 1B, S4A**). We could not test the last two CNVs because patient DNA was no longer available.

One of the focally deleted lncRNA was NR_033883 (also known as or LOC339529). This lncRNA locus is disrupted by two focal CNVs in two distinct ASD/ID patients: 990914 and 9900589 (**Fig. S4A**). The NR_033883 locus neighbors the coding genes *ZFP238* (also known as ZBTB18, ZNF238, and RP58) and *AKT3*. Because the human NR_033883 locus is most proximal but does not overlap *ZFP238*, we propose to refer to this lncRNA as *lnc-ZFP238*. Intriguingly, we previously identified ZFP238 as a key downstream target of the Ascl1 network during MEF to iN cell reprogramming ^11^. Additionally, ZFP238 has an important role in neuronal differentiation during brain development ^28-30^, and thus, *lnc-ZFP238* could have a promising neurogenic role given its high expression pattern during the early stages of direct neuronal reprogramming, as well as in postnatal mouse and human brain (**Fig. S2**).

Another focally deleted lncRNA was the locus harboring human FLJ42709 (or NR2F1-AS1) (**Fig. 1B**) which is adjacent to the protein-coding gene *NR2F1* (also known as *COUP-TF1*), encoding a transcription factor involved in neurogenesis and patterning ^7,31-38^. This lncRNA was previously annotated as “NR2F1-antisense 1” (NR2F1-AS1). However, our RNA-seq analysis showed that at least one isoform of the mouse lncRNA and all detected isoform of the human lncRNA are transcribed divergently from *NR2F1* without antisense overlap ^7^. For scientific accuracy, we therefore propose the name *lnc-NR2F1*. We first asked whether the coding gene *NR2F1* could also be affected by the focal CNVs in ASD/ID patients. Detailed statistical analysis of the primary data taking into account the relative probe density suggested that inclusion of *NR2F1* is not statistically significant compared to the control group. Moreover, we precisely mapped the independent focal deletion found in patient 9900850 by CGH analysis and found only the *lnc-NR2F1* locus to be disrupted (**Fig. 1B**). These results implicated the genetic disruption of *lnc-NR2F1* as likely contributor to complex neurodevelopmental disorders. Chromosomal aberrations encompassing the *lnc-NR2F1* locus and additional genes on chromosome 5q14 have been previously reported in several patients with neurodevelopmental deficits and congenital abnormalities (**Fig. S4B and Table 1**) ^39-42^. However, given that several genes are affected by the deletions, the particular contribution of each gene was difficult to resolve (**Supplementary Text 2**). Across those patients with structural variation encompassing the *lnc-NR2F1* locus in the literature, the minimal deleted region is approximately 230kb, a small area encompassing the genes *NR2F1* and *lnc-NR2F1* (**Fig. S4B and Table 1**). The most notable overlapping phenotype consists of global developmental delay, facial dysmorphism, and hearing loss. Hypotonia and opththalmological abnormalities are also common diagnoses ^39^ (**Fig. S4B and Table 1**). Phenotypic heterogeneity amongst patients could be the result of dosage sensitive genes, polymorphisms on the unaffected allele, genomic variability, gender, and age, amongst others.

Independent of the patients previously reported ^39-42^, we identified a paternally inherited balanced translocation t(5;12)(q15;q15) in a 2 year and 7-month-old male patient (CMS12200) (**Fig. 1C-D and S4B**). Patient CMS12200 was diagnosed with developmental delay, speech delay, significant expressive language delay, and congenital infantile left eye esotropia (**Fig. 1C**). Physical examination revealed small head size (head circumference 48.5 cm; 5^th^-10^th^ centile), and mild fifth finger clinodactyly bilaterally. Other physical features were normal. The patient’s father was diagnosed with dyslexia and stutters, and carried the identical t(5;12) translocation. The patient’s mother had a normal 46, XX karyotype (**Fig. 1C**). Fluorescence *in situ* hybridization and whole genome sequencing defined the chromosomal breakpoints with high precision and revealed that only the *lnc-NR2F1* gene is disrupted in this patient (**Fig. 1D and 1E, S4C**). In humans, three predominant isoforms of *lnc-NR2F1* have been detected in neuronal tissue. The long isoforms (1 and 2) are affected by the chromosomal break, while the gene structure of the short isoform (3) could remain unaffected based on the location of the break (**Fig. 1D**). Further studies by Sanger sequencing of the 5q15 and 12q15 breakpoint-specific junction fragments showed the identical breakpoints in the patient’s father and revealed a loss of 9 nucleotides at the 5q15 chromosome and a loss of 12 nucleotides at the 12q15 chromosome in the patient and his father (**Fig. 1D**). The breakpoint at 12q15 occurred in a coding gene desert and did not disrupt any coding genes, and is predicted to destabilize the affected transcript due to loss of 3’ splice or polyadenylation signals (**Fig. 1D and S4D, S4E**). Importantly, whole genome sequencing data indicated the absence of other deleterious mutations known to be associated with autism or intellectual disability, to our knowledge (**Fig. 1D**). Also, the genes adjacent to the break point (*FAM172A, ARRDC3, KIAA0625, USP15*) are not significantly changed (**Fig. S4E**). Given that *lnc-NR2F1* is the only disrupted gene in this patient family, it is possible that haploinsufficiency is the primary cause for this syndrome and contribute to the phenotypes manifested across patients mentioned above. Further studies including a larger sample size and independent cases are required to conclusively link lnc-*NR2F1* mutations to multiple clinical symptoms described above.

### Molecular and functional characterization of *lnc-Nr2f1*

Given its potential involvement in neurodevelopmental disease, we next sought to investigate the function of *lnc-Nr2f1*. We focused on mouse *lnc-Nr2f1* as experimental approaches are more tractable in mouse models given the availability of a plethora of genetic tools. Remarkably, and in contrast to many other lncRNAs, *lnc-Nr2f1* was not only syntenically conserved in its genomic context (**Fig. S5A**)^43^, but also highly sequence conserved among all human lnc-*NR2F1* isoforms and the three exons in mouse lnc-*Nr2f1* (**Fig. 2A, S5B**). In addition, we identified short stretches of sequence homology (termed microhomology) near the conserved exons (exon 2 and 3 of human lnc-*NR2F1)* across different species, with recurrent sequence motifs and motif order conserved across different species (**Fig. S5C**). All of the above are features hinted at lncRNA functional conservation across different species ^43^.

**Fig. 2.**
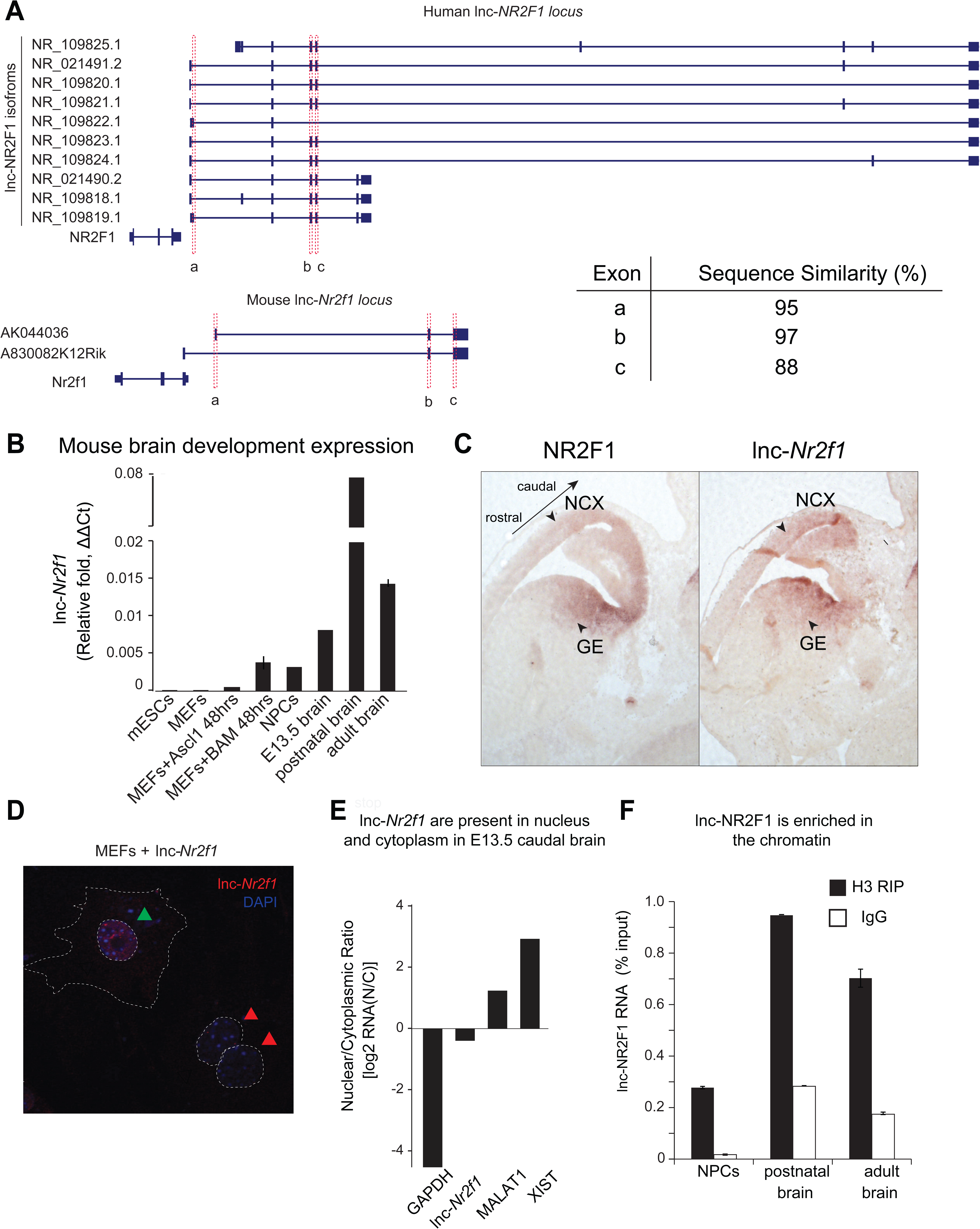
Molecular characterization of mouse *lnc-Nr2f1*. (A) Schematic showing the different isoforms reported by Refseq of the human lnc-*NR2F1* and mouse lnc-*Nr2f1.* Exons highlighted in red are conserved among human and mouse. The table at the bottom right corner shows the sequence similarity as reported by T-COFFEE. The sequence alignment is available as data S7. (B) *Lnc-Nr2f1* expression measured by qRT-PCR across stages of mouse brain development and early stages of iN cell reprogramming. Results show early detection in E13.5 brain, peak expression at postnatal stages, and continued expression through adulthood. (C) In situ hybridization for *Nr2f1* and *lnc-Nr2f1* shows similar expression pattern in E13.5 mouse brain. Highlighted by arrows are neocortex (NCX) and ganglionic eminences (GE) with high expression levels. (D) Cellular localization of *lnc-Nr2f1* by single molecule RNA-FISH in MEFs ectopically expressing the lncRNA 48 hours after dox induction reveals nuclear and cytoplasmic localization, with slight nuclear enrichment. Green arrow points at lnc-*Nr2f1* in the nucleus and red arrows point at the uninfected nuclei. (E) Cellular fractionation of primary neurons derived from E13.5 caudal cortex dissection shows nuclear and cytoplasmic localization of *lnc-Nr2f1*. Chromatin enrichment of *lnc-Nr2f1* by histone H3 RIP-qRT-PCR in brain derived neuronal precursor cells (NPCs), postnatal and adult mouse brain.

Mouse l*nc-Nr2f1* is induced as early as 48 hours after BAM factors are expressed during MEF-to-iN cell conversion and peaks during mid-to-late stages of reprogramming (**Fig. 2B, S2 and S6A)**. In the developing and adult mouse brain, *lnc-Nr2f1* showed a distinct region-specific pattern of expression (**Fig. 2C and S6B**). In the developing telencephalon at E13.5, *in situ* hybridization with a probe against *lnc-Nr2f1* revealed strong expression in the caudolateral part of the mouse cortex and ganglionic eminences (GE), similar to *Nr2f1* expression ^44^ (**Fig. 2C and S6B**).

To determine *lnc-Nr2f1’*s cellular localization, we performed single molecule RNA-FISH in MEFs ectopically expressing *lnc-Nr2f1*, which revealed a nuclear and cytoplasmic but predominantly nuclear localization (**Fig. 2D and S6C**). Consistently, cellular fractionation from primary neurons dissected from caudal region of the cortex showed endogenous localization of *lnc-Nr2f1* in both nuclear and cytoplasmic fractions (**Fig. 2E**). Within the nuclear fraction, *lnc-Nr2f1* is enriched in chromatin as assayed by histone H3 RNA Immunoprecipitation followed by qRT-PCR (histone H3 RIP-qRT-PCR) in brain-derived NPCs, postnatal and adult mouse brain (**Fig. 2F**).

We next wanted to explore potential functional roles of *lnc-Nr2f1* and assessed its role during neuronal induction (**Fig. S7**). We therefore co-expressed *lnc-Nr2f1* (NR_045195.1 or A830082K12Rik) with Ascl1 and asked whether it could promote neuronal conversion over Ascl1 alone as previously observed with other transcription factors (Brn2, Myt1l) ^45,46^. To that end we infected MEFs with Ascl1 with or without *lnc-Nr2f1*, and determined the ratio of TauEGFP-positive cells with neuronal processes over the total number of TauEGFP-positive cells at day 7. We chose day 7 to perform the experiment as it is an early time point for reprogramming. Indeed, the addition of *lnc-Nr2f1* showed an approximately 50% (1.5-fold) significant increase in the number of TauEGFP positive cells with neurites relative to Ascl1 alone (**Fig. S7A and S7B**). This surprising morphological maturation phenotype were only previously observed only with co-expression of transcription factors (Brn2 and Myt1l) with Ascl1 demonstrating a role of *lnc-Nr2f1* in neuronal morphological maturation^47^.

RNA-seq in sorted 7d MEF-iN cells expressing Ascl1, with and without co-expression of *lnc-Nr2f1*, revealed 343 genes significantly changed expression between data sets (RPKM>1, FDR corrected p <0.001) (**Fig. S7C and S7E**). The vast majority of these genes were induced in expression upon *lnc-Nr2f1* expression, with 311 genes up- and 32 down-regulated, suggesting *lnc-Nr2f1* may positively enhance transcription. Gene ontology (GO) term enrichment of up regulated genes showed significant enrichment in biological functions related to plasma membrane (*extracellular region, cell adhesion,* and *transmembrane receptor tyrosine kinase activity*) and neuronal function (*neuron projection, calcium binding, neuron differentiation,* and *axonogenesis*) (**Fig. S7D**). These pathways are consistent with phenotype observed for *lnc-Nr2f1* during iN cell reprogramming of promoting precocious maturation programs (**Fig. S7D**). Amongst the up-regulated genes are well-characterized neuronal and axon guidance genes such as *NeuroD1, Gap43, Tubb4a, Ntf3, Nlgn3, Efnb3, Ntrk3, Bmp4, Sema3d, Slc35d3, Ror1, Ror2, Fgf7*. Additionally, genes previously associated with neurological disorders were similarly up regulated, such as *Mdga2, Clu, Epha3, Chl1, Cntn4, Cdh23*, and *Pard3b* ^48^.

### *Lnc-Nr2f1* **is required for proper neuronal gene expression**

To investigate the contribution of *lnc-Nr2f1* to overall gene regulation, we sought to achieve *lnc-Nr2f1* gain and loss-of-function in one experimental system. We reasoned that mouse ES cells were the best way to accomplish loss-of-function. Since the functional domains of *lnc-Nr2f1* RNA are unknown, non-coding sequences cannot be turned into missense information by frame-shift mutations. Also, a large deletion encompassing the entire 20kb *lnc-Nr2f1* locus may also inactivate interweaved intronic regulatory elements and chromatin structure. Instead, we chose to insert a polyA transcriptional termination signal to eliminate *lnc-Nr2f1* transcripts. We obtained mouse ES cells that were previously genetically characterized in a genome-saturating haploid ES cell mutagenesis screen^49^. One of those ES cell clones had the mutagenesis cassette containing an inverted (therefore inactive) splicing acceptor and polyA site inserted after the first exon of *lnc-Nr2f1* (“Control” thereafter, **Fig. 3A**). The mutagenesis cassette was designed to be conditionally reversible as it is flanked by combinations of loxP sites.

**Fig. 3.**
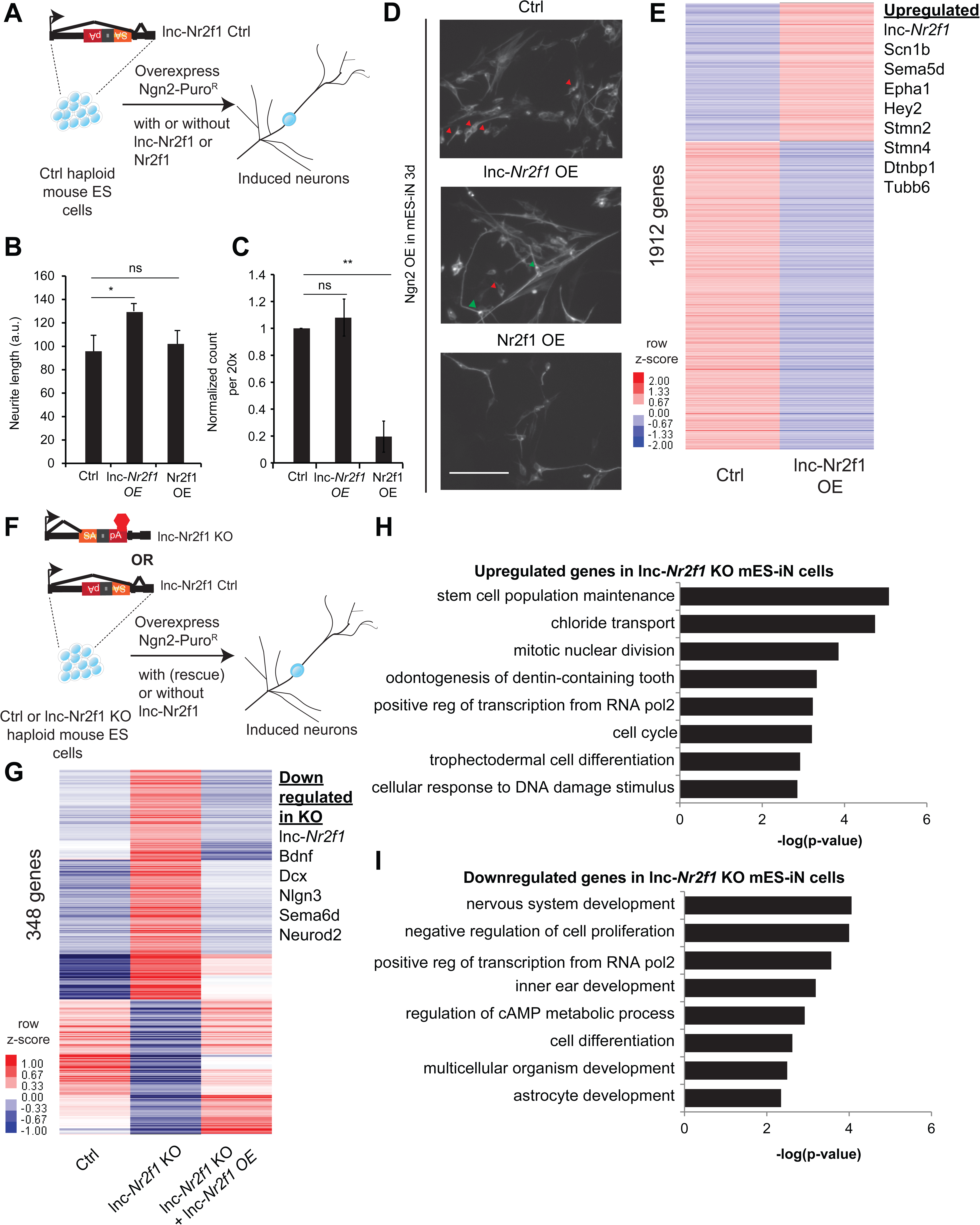
Mouse *Lnc-Nr2f1KO* reveals lnc-*Nr2f1* regulates neuronal genes. (A) Schematic showing the experimental strategy for lnc-*Nr2f1* overexpression. In control mouse ES cells, an inverted construct with a splice acceptor (marked in yellow) and a polyadenylation signal (marked in red) are added after the first exon of the *lnc-Nr2f1.* The mouse ES were infected with rtTA and Ngn2-T2A-puro and mES derived induced neurons (mES-iN) were assayed after 3 or 4 days after dox induction. (B) Graph showing that overexpression of *lnc-Nr2f1* increased the neurite length in day 3 Ngn2 mouse ES derived iN cells relative to the Ctrl. The same effect was not seen with *Nr2f1* overexpression. For each replicate, the individual neurite length for all neurons in each of the five 20x field was manually traced in Fiji. The sequence used for mouse *lnc-Nr2f1* overexpression is available in the supplementary document (n=3, Student t-test, Two-tailed, * indicates p<0.05). Error bars show s.e.m. (C) Graph showing that overexpression of *Nr2f1* decreased the neurite number in day 3 Ngn2 mouse ES-iN cells relative to the Ctrl. The same effect was not seen with *lnc-Nr2f1* overexpression. (n=3, 10 field per replicate, Student t-test, Two-tailed, ** indicates p<0.01). Error bars show s.e.m. (D) β-III-tubulin staining of the day 3 Ngn2 mouse ES derived iN cells for Ctrl, *lnc-Nr2f1* overexpression and Nr2f1 overexpression. Scale bar = 50µm. Red arrow pointed at immature induced neuronal cells with short projection. Green arrow pointed at mature induced neuronal cells with longer projection. Note that the lnc-*Nr2f1* overexpression condition have more mature induced neuronal cells. (E) Hierarchical clustering heatmap of day 4 Ngn2 ES-iN cells between control and *lnc-Nr2f1* overexpression (OE). There are 1912 genes differentially expressed (n=2, FDR corrected p<0.10, Fold change>1.5 fold). Listed to the right are genes which are upregulated upon *lnc-Nr2f1* overexpression. (F) Schematic showing the knocking out strategy for *lnc-Nr2f1*. The *lnc-Nr2f1* knockout mouse ES cells are generated after Cre recombinase introduction to the Ctrl line in Fig.4A. The mouse ES were infected with rtTA and Ngn2-T2A-puro and mES derived induced neurons were assayed after 3 or 4 days after dox induction. (G) Hierarchical clustering heatmap of day 4 Ngn2 ES-iN cells between wild type, *lnc-Nr2f1* knockout (KO) and *lnc-Nr2f1* knockout with *lnc-*Nr2f1 overexpression (OE). There are 348 genes differentially expressed and can be subsequently rescued with *lnc-*Nr2f1 overexpression (n=2, FDR corrected p<0.10). Listed to the right are genes which are upregulated upon *lnc-Nr2f1* KO. (H) Gene ontology of the upregulated genes in *lnc-Nr2f1* knockout day 4 Ngn2 mouse ES-iN cells as compared to the Ctrl. (I) Gene ontology of the downregulated genes in *lnc-Nr2f1* knockout day 4 Ngn2 mouse ES-iN cells as compared to the Ctrl.

To achieve gain-of-function in the same cell system, we first overexpressed *lnc-Nr2f1* in mES cells together with the proneural Ngn2 because that was shown to efficiently and rapidly induce neurons from ES cells ^50^. Again, we quantified the neurite length and neuron count, to test for its effect on increasing maturation kinetics. Consistent with the fibroblast reprogramming results, we saw a significant increase in neurite length upon *lnc-Nr2f1* overexpression though the number of neurons remained the same (**Fig. 3B-D**). In contrast, overexpression of the coding gene *Nr2f1* did not induce these phenotypes, and instead caused a drastic reduction in the number of neurons (**Fig. 3B-D**). These divergent results suggest that *lnc-Nr2f1* functions independently of *Nr2f1*. RNA-seq analysis showed lnc-*Nr2f1* overexpression led to the induction of genes with functions in axon guidance (*Sema5d, Epha1)* and neuronal projection development (*Tubb6, Stmn2, Dtnbp1*) (**Fig. 3E**), confirming the cell biology phenotype (FDR corrected p <0.10, fold change > 1.5).

Next we turned to inactivate *lnc-Nr2f1* function in mouse ES cells. To generate an isogenic knockout line, we treated the conditionally mutant ES cell line with Cre recombinase, which resulted in an inversion of the polyA cassette, which in turn terminates *lnc-Nr2f1* transcription (“*lnc-Nr2f1* KO” thereafter) (**Fig. 3F**). Since *lnc-Nr2f1* is not expressed in ES cells, we differentiated *lnc-Nr2f1* KO and control mES cells into induced neuronal cells by Ngn2 overexpression as above to assess the transcriptional consequences of loss of *lnc-Nr2f1* in a neuronal context. RNA-seq showed that 348 genes were differentially expressed between the control and the *lnc-Nr2f1* KO neurons, which can be subsequently rescued with *lnc-Nr2f1* overexpression (FDR corrected p <0.10) (**Fig. 3G and Fig. S7F**). Consistent with target genes in our *lnc-Nr2f1* overexpression study, we found *lnc-Nr2f1* KO led to down regulation of neuronal pathfinding and axon guidance genes such as *Sema6d* and proneural bHLH transcription factor *Neurod2* as well as deregulation of genes associated with autism spectrum disorder such as *Bdnf, Dcx and Nlgn3* ^48^ (**Fig. 3G**). The transcriptional abnormalities were reversed by enforced expression of *lnc-Nr2f1* from a heterologous construct via lentiviral transduction. The rescue data indicate that the downregulation of neuronal genes and the upregulation of ectopic genes are caused by the loss of *lnc-Nr2f1* expression in the knock out cells and unlikely by disruption of the nearby DNA regulatory elements due to the insertion of the targeting cassette. Gene Ontology analysis of the downregulated genes in *lnc-Nr2f1* KO neurons revealed enrichment for terms related to neural functions (*regionalization, central nervous system development and neural precursor cell proliferation*), whereas the upregulated genes are enriched in development of non-neuronal tissues such as *circulatory system and skin development* (**Fig. 3H and 3I**). Finally, the rescued genes overlap significantly with curated autism risk genes by Basu et. al. ^48^ (p=0.0012, Chisquare) (**Fig. S7G**).

To further distinguish the function of *lnc-Nr2f1* vs *Nrf1*, we generated *Nr2f1* heterozygous and homozygous KO mouse ES cell lines using CRISPR/Cas9 (**Fig. S8A**). When we performed qRT-PCR on the day 4 iN cells generated from the Ctrl, *Nr2f1* heterozygous and homozygous null mES cells, we found that the level of both *Nr2f1* and *lnc-Nr2f1* RNA transcripts did not change (**Fig. S8B**), However, protein quantitation confirmed that Nr2f1 protein level was reduced or eliminated in the heterozygous lines (clone 21 and 44) or homozygous null clones (Clone 2, 11 and 18), respectively (**Fig. S8C**). The *Nr2f1* KO did not affect *lnc-Nr2f1* expression (**Fig. S8B**) and had no impact on the neurite length or number in mES-iN cells (**Fig. S8D and S8E**). In summary, both gain and loss of function studies demonstrated that *lnc-Nr2f1* plays a role in the transcriptional regulation of a gene network involved in neuronal maturation pathways that ultimately resulted in faster acquisition of a mature neuronal identity in both MEFs and mES cells and is functionally distinct from its neighboring coding gene, *Nr2f1*.

### Mouse *lnc-Nr2f1* **binds to distinct genomic loci regulating neuronal genes**

As described above, histone pull-down experiments suggested an association of lncNr2f1 with chromatin. We therefore sought to map the precise lnc-*Nr2f1* genome wide occupancy and performed Chromatin Isolation by RNA precipitation followed by sequencing (ChIRP-seq) on day 4 mES-iN (**Fig. 4A, S9A-B**). To minimize background sequencing we used even and odd probes targeting *lnc-Nr2f1* in replicate experiments and only considered the overlapping peaks from both experiments (**Fig. 4B**). Both even and odd probes pulled down *lnc-Nr2f1* efficiently. There are 14975 peaks called by MACS, and the peak signals are consistent between replicates (n=4) with little signal in RNase-treated control samples (**Fig. 4B**). We obtained 1092 high confidence peaks with further filtering for the most significant and reproducible binding events (see methods for filtering criteria). As an example, *lnc-Nr2f1* binds to the intronic region of *Nrp2*, a gene with known roles in neuronal pathfinding (**Fig. 4C**). To understand which gene ontology terms are enriched in the genes adjacent to the mES-iN ChIRP peaks, we performed GREAT analysis and associated the 1092 *lnc-Nr2f1* binding sites to 1534 genes. GO term analysis revealed that these genes that enriched for neuronal terms such as *central nervous system development, synapse organization and chemical synaptic transmission* (**Fig. 4D**). Using the publicly available ChIP-seq peak sets for CTCF, enhancer, H3K27ac, H3K4me3 and PolII obtained from mouse adult cortex and E14.5 brain, we found significant enrichments of those peaks co-localizing with the enhancers, H3K27ac and H3K4me3 marks relatively to the background (**Fig. 4E and S9D**) ^51^. DNA motif analysis of *lnc-Nr2f1* binding sites revealed several basic helix loop helix (bHLH) factor motifs that are significantly enriched (*NeuroD1, Atoh1, Olig2 and Ptf1a*) (**Fig. S9C**). Finally, to understand whether the *lnc-Nr2f1* regulates the genes listed in figure 3E, we overlapped the 1534 genes adjacent to the mouse *lnc-Nr2f1* ChIRP peaks with the genes up or downregulated upon *lnc-Nr2f1* overexpression, we found 177 common genes between the two lists (p<0.0001, Chi Square), suggesting that those genes might be direct lnc-Nr2f1 targets since they are occupied by lnc-Nr2f1 RNA and are significantly altered when manipulating lnc-Nr2f1 (**Fig. S9E**)

**Fig. 4.**
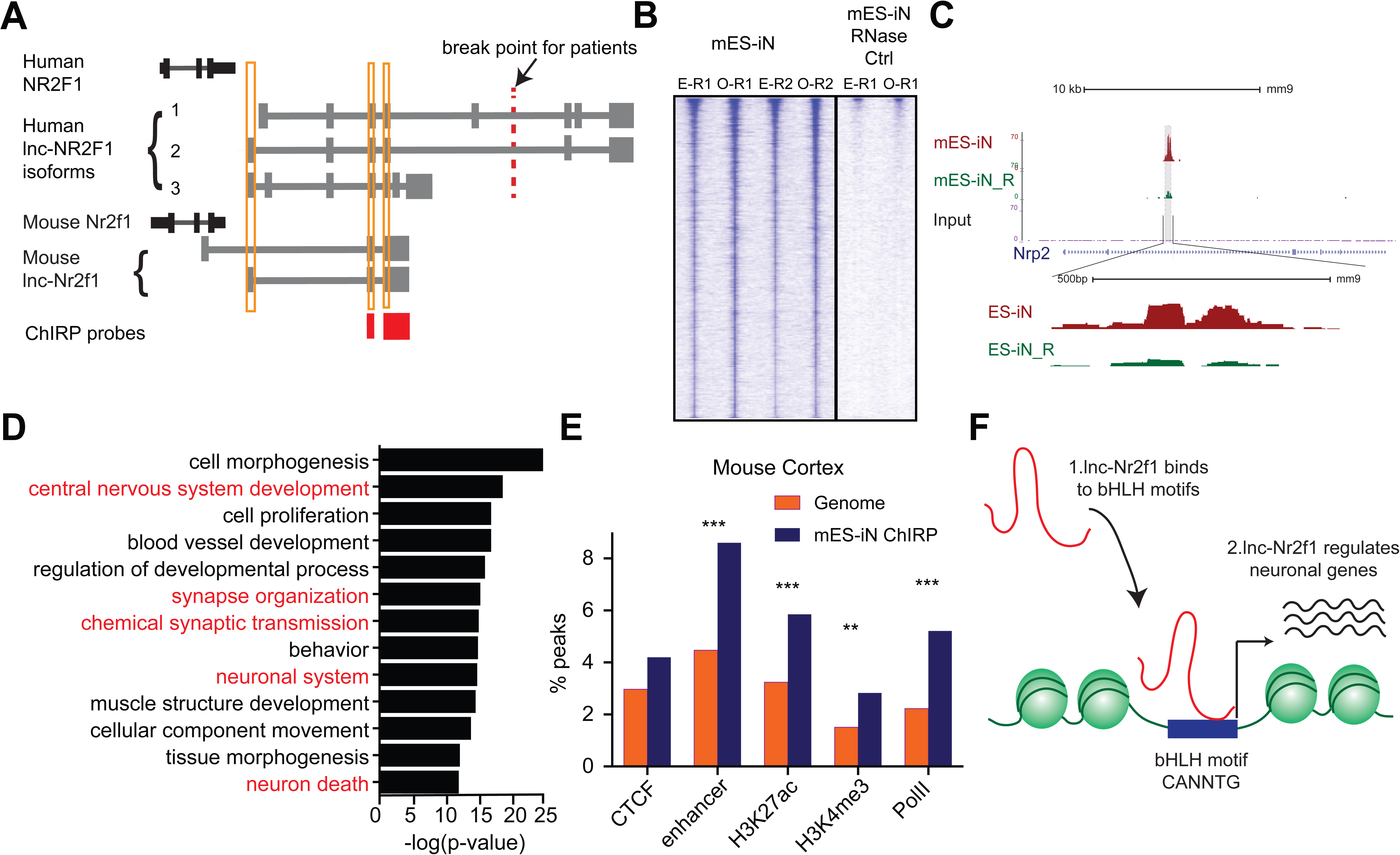
*lnc-Nr2f1* binds to distinct genomic loci regulating neuronal genes. (A) Schematic showing the location of ChIRP probe for mouse *lnc-Nr2f1* (highlighted in red). Yellow lines represent the conserved exons between mouse and human lnc-*Nr2f1*. (B) Heatmaps representing genome-wide occupancy profile for mouse *lnc-Nr2f1* in day 4 Ngn2 mouse ES-iN cells and the RNase control obtained by ChIRP. There are 14975 significant peaks called with respect to the RNase treated control. E and O represents even and odd probes respectively. (C) UCSC browser track showing the binding site within the intronic region of Nrp2. The “R” represents the RNase treated control. (D) Gene ontology terms associated with genes adjacent to the high confident mES-iN ChIRP-seq peaks. Terms highlighted in red are terms related to nervous system development. (E) Percentage of mES-iN ChIRP-seq peaks which overlap with CTCF, enhancer, H3K27ac, H3K4Me3 and PolII defined in mouse cortex relative to the control. (*** represents p<0.0001, ** represents p<0.01, Chi-square test) (F) Proposal mechanism of lnc-*Nr2f1* action. lnc-*Nr2f1* binds to the genomic region enriched with bHLH motif and regulates the downstream neuronal genes.

### Human *lnc-NR2F1* **shows isoform-specific chromatin binding**

The balanced chromosomal translocation t(5;12) detected in patient CMS12200 disrupts the long *lnc-NR2F1* while the short isoform appears unaffected. We therefore hypothesized that the *lnc-NR2F1* might have isoform-specific functions and the long isoforms are contributing to the phenotype observed in patient CMS12200. Given that *lnc-NR2F1* is highly expressed in human brain tissue and it has high sequence conservation between mouse and human (**Fig. 5A**), we next sought to determine whether human *lnc-NR2F1* had a similar role in neuronal reprogramming as the mouse transcript and whether the different isoforms may have distinct functions. Therefore, we individually expressed each of the three human *lnc-NR2F1* isoforms in MEFs, and measured their ability to enhance Ascl1-mediated neuronal reprogramming, as judged by morphological complexity of TauEGFP cells (**Fig. 5B**). The long human *lnc-NR2F1* isoform 2 significantly increased the proportion of TauEGFP cells with projections, albeit with a slight smaller magnitude than the mouse lncRNA. Intriguingly, long isoform 1 inhibited neuronal maturation while the short isoform 3 had no significant effect (**Fig. 5B**). Thus, different isoforms of *lnc-NR2F1* may possess differential regulatory activity. Long *lnc-NR2F1* isoforms disrupted by chromosomal translocation in patient CMS12200 can impact neuronal maturation, while the short *lnc-NR2F1* isoform remaining intact in patient CMS12200 did not have a detectable effect on neuronal complexity.

**Fig. 5.**
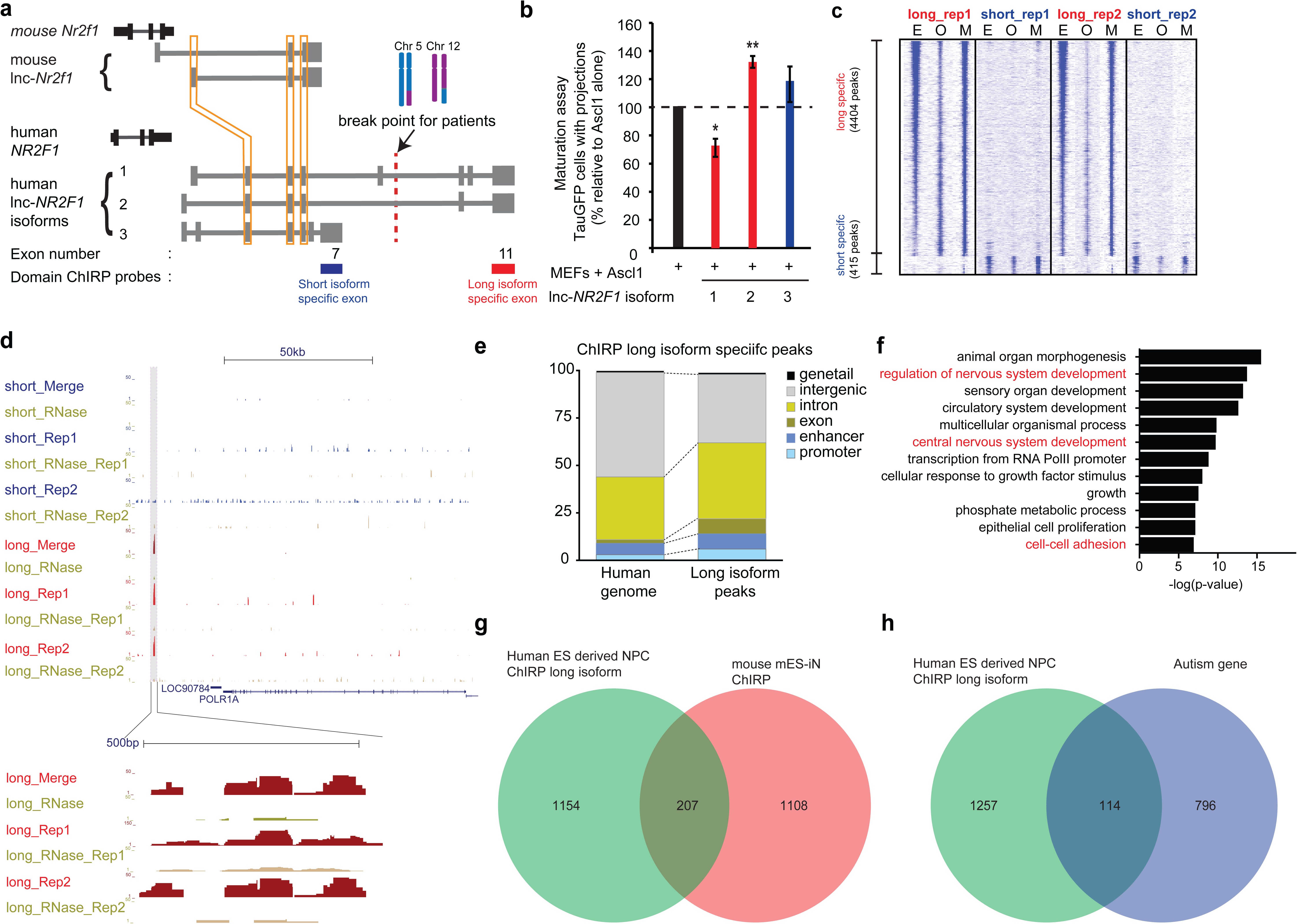
Human *lnc-NR2F1* shows isoform-specific chromatin binding. (A) Schematic showing the location of the ChIRP probes target the short isoform-specific exon (exon 7) and long isoform-specific exon (exon 11). The red line denotes the break point for the patient. (B) Overexpression of human *lnc-NR2F1* isoforms in combination with Ascl1 relative to MEFs expressing Ascl1 alone. The graph quantifies the proportion of TauGFP positive cells with projections normalized to number of TauGFP cells. TauGFP cells with projections longer than three times the diameter of the cell body were counted and normalized to the total number of TauGFP positive cells. The sequences for human *lnc-Nr2f1* isoforms is available in the supplementary documents (n=3, Student t-test, two tailed, scale bar= * represents p<0.05, ** represents p <0.01). Error bars show s.e.m. (C) Heatmap representing genome-wide occupancy profile for domain ChIRP performed using probes specific to the long and short isoform-specific exon of *lnc-NR2F1* in human ES derived neural progenitor cells (NPC). There are 4404 and 415 significant peaks called relative to the RNase control for the long and short isoform respectively. E, O and M represents even, odd and merge track respectively. (D) UCSC browser track showing the site within the promoter region of LOC90784 bound by the long isoform-specific exon (exon 11) but not the short isoform-specific exon (exon 7) (E) Bar graph showing the distribution of the 913 high confident long isoform-specific peaks. The long isoform-specific peaks are enriched in the introns, exons, promoters and enhancers but depleted in the intergenic regions. (F)Gene ontology terms associated with genes adjacent to the human ES derived NPC ChIRP-seq high confident peaks. Terms highlighted in red are terms related to nervous system development. (G) Venn diagram representing the peak associated gene overlap between the domain ChIRP of the long isoform-specific exon (exon 11) from human ES derived NPC and mouse mES-iN ChIRP. (p<0.0001, Chi square test) (H) Venn diagram representing overlap between genes involved in the autism risk and genes identified by the domain ChIRP of the long isoform-specific exon (exon 11) from human ES derived NPC. (p<0.0001, Chi square test).

Due to *lnc-NR2F1*’s strong association with chromatin and isoform-specific function, we hypothesized that different domains of *lnc-NR2F1* may have differential chromatin localization. To test this idea, we mapped the genome-wide localization of different RNA domains in *lnc-NR2F1* using domain-specific Chromatin Isolation by RNA precipitation followed by sequencing (domain ChIRP-seq) ^51,52^. We performed *lnc-NR2F1* ChIRP-seq in human neural progenitor cell (hNPC) differentiated 12d from human embryonic stem cells using dual SMAD inhibition protocol ^53^ (**Fig. S10A**). We performed ChIRP-seq separately with two orthogonal probe sets (termed odd and even sets) against two different domains of *lnc-NR2F1* (long isoform-specific exon 11 and the short isoform-specific exon 7) and only accepted concordant results between the odd and even probe sets. There are approximately 10-fold more genomic occupancy for the long vs. short isoform of *lnc-NR2F1*: 4404 ChIRP-seq peaks for exon 11 (n=4) vs. 415 ChIRP-seq peaks for exon 7 (n=4), respectively. It is unlikely that low expression or inefficient pulldown of the short isoform are the cause of the difference given that we detected comparable level of long and short isoform in hNPC and obtained similar recovery of the long and short RNA isoform (**Figure S10B and S10C**). Consistent with our hypothesis of domain specific chromatin localization, genomic occupancy sites of different *lnc-NR2F1* exons showed limited overlap of peaks (**Figure 5C**), suggesting that the long- and short-specific exon might be each binding to different genomic loci and regulating different subsets of downstream genes. For example, only the long isoform-specific exon has a distinct binding site surrounding *POLR1A* (**Figure 5D**).

We then assessed the transcriptional response which the three isoforms of human *lnc-NR2F1* varied quantitatively: The shortest isoform, human *lnc-NR2F1* isoform 3, had the lowest number of differentially regulated genes (5 downregulated and 4 upregulated) compared to isoform 1 (1147 downregulated and 414 upregulated) and isoform 2 (141 downregulated and 45 upregulated) (**Figure S10F**). This is consistent with our hypothesis that the long isoform is the functional one in neurogenesis and in patients with neurodevelopmental disorders including ASD.

After further filtering the peaks for high confidence peaks, we obtained 913 high confidence peaks for long-isoform domain ChIRP and no peaks for short-isoform domain ChIRP (see methods for filtering criteria). To further characterize *lnc-NR2F1* occupancy patterns, high confidence ChIRP peaks were classified according to distance to putative *cis*-genes (**Figure 5E and S10D**). Relative to the human genome, the long isoform-specific peaks are more enriched in the exonic, intronic, enhancer and promoter regions and depleted in the intergenic regions (**Figure 5E**). Furthermore, ChIRP peaks are characterized by chromatin state model, which defines human genome with 25 chromatin states using 12 biochemical features (histone modifications, DNA accessibility, DNA methylation, RNA-seq and other epigenetic signals) ^54^. *Lnc-NR2F1* ChIRP peaks are enriched in promoter, enhancer, and transcribed regions, compared to the whole genome which majority is in the quiescent state (**Figure S10D**).

The 1361 high confidence peak associated genes by the long isoform-specific domain ChIRP are all enriched for neuronal specific biological terms such *central nervous system development, cell-cell adhesion* and *regulation of nervous system development* (**Figure 5F**). There is also a significant overlap between the genes adjacent to peaks from the long isoform-specific domain ChIRP and the genes adjacent to the mouse *lnc-Nr2f1* ChIRP, indicating a possible conserved functions of *lnc-Nr2f1* between mouse and human (207 genes overlapped, Chi-square test, p<0.0001) (**Figure 5G**). We also observed a significant overlap between the genes adjacent to the peaks from the long isoform-specific domain ChIRP and the autism risk genes (114 genes, Chi-square test, p<0.0001) (**Figure 5H**). Notably, peaks of long isoform-specific exon 11 ChIRP are enriched for multiple basic-helix-loop-helix motifs (Ascl1, NeuroD1, Olig2 and Atoh1) which all share the CANNTG motif ^55^(**Figure S10E**). Mouse ES-iN cell ChIRP peaks are also enriched for similar motifs suggesting possibly conserved mechanisms of *lnc-NR2F1* in human and mouse. Given the pervasive roles of bHLH proteins in neuronal development and in induced neuronal reprogramming, the binding preference of *lnc-* NR2F1 suggests a biochemical basis for the functional cooperativity with proneural bHLH factors (Ngn2 and Ascl1). In summary, we conclude that the *lnc-NR2F1* isoforms have different genomic occupancy and transcriptional effects. The long isoform showed most biological activity and chromatin binding and is also solely affected in patient CMS12200.

## Discussion

Given the stringency required to rewire a cellular state from an unrelated lineage, factors expressed during direct reprogramming likely have an active role in the establishment of the new cell identity. Direct neuronal lineage induction represents a synchronized and streamlined conversion of cell fate, and should be a powerful system to enrich for lineage-specific regulatory factors. In this study, direct conversion of fibroblasts to induced neuronal cells enabled the identification of lncRNAs with unique properties in establishing neuronal fate via neurogenesis or maturation programs. The pipeline described in this study may be extrapolated to identify potential lncRNA regulators of other specific cellular states.

Because the brain is the organ with the greatest number of unique cell-type specific lncRNAs ^56^, our approach may be useful to identify lncRNAs with roles in neural lineage specification. Indeed, we identified *lnc-Nr2f1* as functional player in neuronal maturation and pathfinding. Remarkably its sequence is remarkably conserved between the first few exons of mouse and human *lnc-NR2F1* which is an atypical pattern for lncRNAs ^57,58^. Consequently, we found that there is high synteny, sequence and microdomain conservation between mouse and human *lnc-NR2F1*. These observations suggest that some lncRNAs may have been functionally conserved throughout evolution.

In this study, we focus on the functional characterization o f *lnc-NR2F1* locus because it is recurrently mutated in human patients with ASD/ID. We identified a patient, whose genome harbors a balanced translocation disrupting the *lnc-NR2F1* locus without any other detectable pathogenic genetic variations and shows abnormal neurodevelopmental symptoms; therefore, implicating this lncRNA as a critical regulator of brain development and function. The father of the proband carries the same translocation and suffers from dyslexia and stuttering, suggesting that the phenotype may be transmitted in a Mendelian manner. However, the much milder phenotype of the father implies that additional genes, environmental factors, or compensatory neuronal circuitry acquired during adulthood may influence the severity of the outcome.

Given the close genomic proximity of *lnc-NR2F1* and *NR2F1* increased attention must be devoted to consider the possibility of a contribution of the coding gene. Since unknown regulatory elements for the coding gene could be affected the human genetics data are not decisive. However, several functional experiments point to a contribution of *lnc-NR2F1* rather than the coding gene. First, gain and loss of function studies as well as chromatin localization clearly show that *lnc-NR2F1* acts in trans to affect gene expression. Second, we can rescue the phenotype of the *lnc-Nr2f1* KO by overexpression of *lnc-Nr2f1* mRNA. Third, *lnc-Nr2f1* overexpression in *lnc-Nr2f1* KO cells does not affect *Nr2f1* expression. Fourth, ChIRP-sequencing of *lnc-Nr2f1* in induced neurons derived from mES cells does not show binding of *lnc-Nr2f1* in the *Nr2f1* promoter region (**Fig. 4F**). The only definitive answer may be obtained from human postmortem tissue analysis of affected patients.

Neurodevelopmental and neuropsychiatric disorders are complex diseases manifesting in a spectrum of phenotypes. We integrated lncRNA expression pattern, *in vitro* functional screen, and human genetic data to pinpoint potentially causal lncRNAs. We concentrated on genomic lesions affecting lncRNAs, which have been largely understudied regulatory factors in these diseases, and connected them to specific phenotypes. We found several of the lncRNA candidates were disrupted by focal chromosomal aberrations in patients diagnosed with ASD/ID, establishing a link between human disease and lncRNA function. The advent of next generation sequencing has greatly improved the ability to pinpoint causal disease mutations in protein coding genes including the discovery of novel autism genes. Most of the other functional regions of the genome, however, have largely been ignored as part of exome sequencing approaches. While full genome sequencing of patients is beginning, functional interpretation remains a daunting challenge. We present a strategy to begin to characterize the functionally important non-coding regions as it relates to disease. Our work highlights lncRNA mutations as an understudied and important potential next frontier in human genetics related to neurodevelopmental disease.

## Acknowledgements

We thank Cindy Skinner, Mrs. Sydney Ladd, Dr. Barbra R. DuPont, Dr. Katie R. Clarkson for patient recruitment and evaluation and members of our labs for discussion and advice. We thank the Stanford Functional Genomic Facility especially Vanita for her assistant in the project. This project is supported by NIH RC4-NS073015 (H.Y.C., M.W.), P50-HG007735 (H.Y.C.), California Institute for Regenerative Medicine (M.W., H.Y.C.), NIH R01 HD39331 (A.K.S.) and Self Regional Healthcare Foundation Funds (A.K.S.). C.E.A. was supported by California Institute of Regenerative Medicine Training Grant and Siebel Foundation. O.L.W. was supported by a NSF fellowship. M.W. is a NYSCF–Robertson Stem Cell Investigator. Haplobank is generously funded by Nestlé Institute of Health Science NIHS as well as the Austrian National Bank (OeNB) and Era of Hope/National Coalition against Breast Cancer/DoD. U.E. is supported by the Austrian Academy of Sciences, the Austrian National Bank (OeNB), and is a Wittgenstein Prize fellow. J.M.P. is supported by an Advanced ERC grant and an Era of Hope/DoD grant. E.E. is an Investigator of the Howard Hughes Medical Institute.

## Author contributions

C.E.A., O.L.W., Q.M., M.W. and H.C. conceived the project. C.E.A., O.L.W., Q.M., M.W. and H.C. wrote the paper with inputs from all authors. C.E.A., and O.L.W. conducted lncRNA discovery, reprogramming, and perturbation experiments. M.O. performed the in situ hybridization and cellular fractionation assays. C.E.A and Q.M. performed the *lnc-Nr2f1* perturbation experiment. C.E.A., Q.M., and R.A.F. conducted the ChIRP-seq experiments. C.E.A, O.L.W., B.T.D., Q.M., J.X., Q.Y.L. analyzed RNA sequencing data. B.C. and E.E. analyzed lncRNA loci recurrently mutated in autism cohorts. S.F., L.D-R., L.W., and A.S. characterized the t (5:12) family and human *lnc-NR2F1*. J.X., B.T.D. helped with data analysis.

## Data availability statement

Datasets used in this paper and their corresponding source, the qRT-PCR primer sequences and the ChIRP-sequencing probes can be found in the supplementary documents. The datasets generated during and/or analyzed during the current study are available in the NIH GEO repository (GSE115079)

## Materials and Methods

This method section was organized into four categories: animal and human protocols, cell culture, computational and sequencing methods and biochemistry. Within each category, method descriptions were arranged in the order they appear in figures.

### Animal and human protocols

#### Animal

All mouse work was performed according to IACUC approved protocols at Stanford University. Samples in the paper were obtained without determining their sex. All animals were housed in an animal facility with a 12hr light/dark cycle.

#### Human subjects

Patient study was approved by the Institutional Review Board of the Self Regional Healthcare, Greenwood, SC.

### Cell culture and tissue dissection

#### Cell culture

Mouse embryonic fibroblasts (MEF) were derived from E13.5 Tau::EGFP embryos and cultured in MEF media [500ml of DMEM (Gibco), 50ml of Cosmic Calf Serum (Thermo Scientific), 5ml of Non-essential amino acid, 5ml of Sodium Pyruvate, 5ml of Penicillin/Streptomycin (Gibco), 4ul of β-mercaptoethanol (Sigma)].

Mouse haploid embryonic stem cells were cultured in mouse embryonic stem cell media [341.5ml DMEM (Gibco), 50ml Knockout Serum Replacement (Gibco), 12.5ml of Cosmic Calf Serum (ThermoScientific), 4.2ml of Penicillin/Streptomycin, 4.2ml of Non-essential amino acid, 4.2ml of Sodium Pyruvate (Gibco), 4ul of β-mercaptoethanol (Sigma) with leukemia inhibitory factor, 3µM of CHIR99021 and 1µM of PD3259010 (Both Tocris, Final concentration)].

Human embryonic stem cells (HUES9) were cultured in mTESR media (Stem Cell Technologies). The experiments were performed in accordance with California State Regulations, CIRM Regulations and Stanford’s Policy on Human Embryonic Stem Cell Research.

#### Mouse postnatal/adult brain dissection

Briefly, forebrains were dissected from TauGFP heterozygous E13.5 embryos in cold HBSS. Neural stem cells (NSC) were propagated in DMEM/F12 with N2 and B27 supplements (Invitrogen) with 20ng/ml of FGF2 and 10ng/ml of EGF. Postnatal brains (Postnatal day 0) and adult brains (three weeks old) were obtained from C57BL6 mice. To obtain postnatal brains, pups were anaesthetized in an ice bath before the whole brain was removed. To obtain adult brains, mice were euthanized using cervical dislocation before dissecting the whole brain out. For both adult and postnatal brains, they were manually dissociated to fine pieces before being digested in 0.25% trypsin for 30 minutes. They were triturated from time to time until a clear suspension was obtained. The cells were spun down at 1000rpm for 5 minutes before proceeding to glutardehyde fixation.

#### Reprogramming of mouse fibroblasts to induced neuronal cells (iN cells)

We followed protocols previously described (Wapinski et al., 2013). Briefly, mouse embryonic fibroblasts harvested from E13.5 Tau::EGFP embryos were plated at a density of 25000 cells/cm^2^. The next day, lentiviruses carrying TetO-FUW-ASCL1 and FUW-rtTA were added. Doxycycline (Final concentration: 2µg/ml, Sigma) in MEF media was added to the wells. Media was changed to neuronal media [N2 + B27 + DMEM/F12 (Invitrogen) + 1.6ml Insulin (6.25mg/ml, Sigma)] + doxycycline two days after doxycycline induction. Subsequently, media was changed every three days. To obtain a pure population of day 7 TauEGFP positive neurons, the cells were digested using 0.25% trypsin (Invitrogen) and subjected to FACS. Forward and side scatters were used to exclude doublets and dead cells. The gating for GFP was set with a negative control (MEF).

#### Maturation screen for *lnc-NR2F1*

To examine whether the lncRNA candidates can facilitate mouse embryonic fibroblasts (MEFs) to induced neuronal cells reprogramming, mouse and the three human isoforms for *lnc-NR2F1* were synthesized (sequences in the supplementary documents) and cloned into the TetO-FUW or TetO-PGK-blast^R^ backbone respectively (available from Addgene). To examine whether those lncRNA candidates can help facilitate the maturation process, the number of MAP2 positive neuronal cells with projections three times the diameter of the cell body was counted at day 7 and normalized to the total number of MAP2 positive cells. For neurite length measurement, Simple Neurite Tracer (ImageJ) was used manually to track neurite.

#### Reprogramming of mouse embryonic stem cells to induced neurons

We followed the protocol previously described ^50^. Mouse embryonic stem cells were plated single cell and infected the next day with TetO-NGN2-T2A-PURO^R^ and FUW-rtTA. Doxycycline was added to the wells the next day. To select for only Ngn2 transducing cells, puromycin (Final concentration: 2µg/ml, Sigma) was added in addition to doxycycline the next day and kept for 3 days.

#### Generating *lnc-Nr2f1* KO ES-iN cells

We obtained mouse ES cells that were previously generated in a genome-saturating haploid ES cell mutagenesis screen ^59^. We identified one ES cell clone had the mutagenesis cassette containing a splicing acceptor and polyA site inserted after the first exon of *lnc-Nr2f1*. The orientation of the polyA site is in reverse from the transcription direction of *lnc-Nr2f1* so it’s non-disruptive. The insertion is confirmed by PCR and sequencing. For generating *lnc-Nr2f1* KO clones, we did nucleofection of cre recombinase to invert polyA cassette since the polyA cassette is flanked by loxP sites. After nucleofection, we plated cells at low density and picked single colonies for testing the polyA inversion. The control and KO clones were then expanded for a few passages, allowing majority of them to become diploid cells. The homozygous diploid cells were then plated at 300K cells/6 well in mES media at day 0. They were then infected with TetO-Ngn2-T2a-puro, FUW-rtTA and TetO-GFP the next day. At day 2, the media was changed to neuronal media [N2 + B27 + DMEM/F12 (Invitrogen) + 1.6ml Insulin (6.25mg/ml, Sigma)] and doxycycline (Sigma, Final conc: 2µg/ml) was added. At day 3, puromycin (Sigma, 2ug/ml) was added to the neuronal + dox media. RNA was harvested four days after dox induction.

#### Generating *Nr2f1* KO ES-iN cells

For CRISPR/Cas9 genome editing of Nr2f1, gRNAs targeting second exon of Nr2f1 are cloned to a plasmid (pSpCas9(BB)-2A-Puro, pX459, Addgene #62988) expressing both the Cas9 protein and the gRNA. gRNA sequences were designed using the online tool (http://crispr.mit.edu/) provided by the Zhang lab (gRNA sequence used: CATGTCCGCGGACCGCGTCG). 24-48 hours after ES cell nucleofection, puromycin was added to select for 2-3 days. The remaining cells were plated at plate 100, 300, 1000, 3000 cells per plate for picking single colonies. Genomic DNA of each single colony was extracted using QuickExtract™ DNA Extraction Solution (Epicentre, QE09050). This extract was then used in a PCR of the genomic region that had been targeted for knock out (Fwd primer: AGAGACACCTGGTCCGTGAT. Reverse primer: GAGCCGGTGAAGGTAGATGA). PCR products were then Sanger sequenced to identify clones that would result in frameshifts and truncated Nr2f1. Sequence alignment and genomic PCR primer design was carried out using SnapGene software and cutting efficiency is calculated using web tool TIDE (https://tide-calculator.nki.nl/).

### Computational and sequencing methods

#### LncRNA discovery pipelines (related to Figure S1)

TopHat was used for de novo alignment of paired-end reads for each of the samples. An assembled transcriptome was built from merged time points using Cuffmerge function. The de novo iN transcriptome was compared to RefSeq genes and annotated protein coding genes were removed, while non-coding genes annotated as “NR” were kept. Expression level of genes was calculated in unit of fragments per kilobase of exon model per million mapped fragments (FPKM). Genes with low FPKM (average log2 FPKM across all samples less than 1) were removed. Genes with p-value<0.05 and at least two-fold expression change during iN reprogramming were defined as significant.

#### Histone H3 RIP-seq (related to Figure S1F)

RNA isolated from H3 RIP was amplified and converted to cDNA using Nugen Ovation RNA-Seq System V2. The product was sheared using Covaris to 100-300bp. Libraries were prepared using SPRIworks system for Illumina sequencing. The following antibodies were used for RIP: Rabbit anti-H3 (Abcam ab1791) and rabbit IgG (Abcam ab37415). For the H3 RIP-seq analysis we used a similar pipeline to bulk RNAseq assays. We first remove duplicate reads, clip adaptor sequences, discard short reads. Reads were then aligned to mm9 using Tophat2. Using samtools we convert the files from sam file to bam format. Filtered reads are normalized to sequence depth, and subsequently we calculate RPKM. The RIP-seq experiment was conducted with H3 and IgG antibodies. We sequenced both libraries and also 1% input material. To determine whether a lncRNA is enriched we compute the number of reads from H3 relative to input, and IgG relative to input using an in-house Perl script (<u>rnaexp_rpkm.pl</u>). We then calculated the fold-enrichment between H3 and IgG RIPs. Since the background was very low, anything greater than 2-fold and p<0.05 between H3 and IgG was considered enriched. Experiments in NPCs, MEFs, and adult brain were conducted in biological replicates. Only lncRNAs reproducibly enriched in the H3 RIP from biological replicates were considered chromatin associated and display as binary in the Figure S1F.

#### Co-expression analysis for lncRNAs (related to Figure S1G)

We first obtained mouse RNAseq data from ENCODE, and calculated the RPKMs for all transcripts including coding and non-coding. We then for each non-coding RNA, calculated the Pearson correlation of the non-coding RNA with every coding transcript. If the correlation is greater than 0.5, this non-coding RNA was defined as positively correlated with the coding gene, and if the correlation is less than 0.5, it was defined as negatively correlated. We then obtained a matrix of coding genes versus non-coding genes, with positive (+1) and negative (−1) correlations as values in the matrix. We then use the GeneSets function in Genomica software from (http://genomica.weizmann.ac.il/), and generated a enrichment (−log(p-values)) matrix for iN lncRNAs associated with Gene Ontology terms based on similar expression pattern with mRNAs. The default settings set in the software were used.

#### Overlap with CNV morbidity map (synteny of coordinates, significance calculation) (related to Figure 2A)

To find syntenic conservation from mouse to human, UCSC Genome Browser tool Liftover was used. To determine the potential role of lncRNAs in neurodevelopmental disorders, we analyzed array CGH profiles from 29,085 children with intellectual disability and developmental delay that were submitted to Signature Genomics Laboratories, LLC, for clinical microarray-based CGH. The CNV map intersecting lncRNAs was compared with that of 19,584 healthy controls ^25^(dbVar nstd100). Focal enrichment was calculated using fishers exact test statistics and odds ratios comparing cases and control CNV counts at each locus. Validation of focal CNVs affecting lncRNAs of interest was performed on a custom 8-plex Agilent CGH array using standard methods ^25^.

#### Ingenuity Variant Analysis (related to Figure 2A)

Using Ingenuity Variant Analysis software, we filtered 4,038,671 sequencing variants and obtained a list of 45 variants possibly related with the patient’s phenotype. This list included three structural variants (deletions) and one gene fusion. We verified by Sanger sequencing each of the variants associated with disease and found them to be either false positives or non-causative.

The 16.8 Mb large deletion on Chromosome 11 was found to be false positive based on the observation of heterozygosity in the deleted region in whole genome sequencing data. The same false-positive was also shown in the whole genome sequencing data of other two translocation patients. The gene fusion between *ARHGEF3* on Chromosome 3 and *TRIO* on Chromosome 5 was determined as false positive by Sanger sequencing. One fragment of *ARHGEF3* (132bp) intron sequence was inserted into an intron of *TRIO*. The insertion led to the false detection of gene fusion. RT-PCR proved that mRNA splicing of *TRIO* was not affected by this insertion and qRT-PCR proved that the expression level of *TRIO* was not affected. For the rest of variants, we reviewed the reported functions of genes having these variants and copy number variation information in these regions in Database of Genome Variants. We found seven variants occurred in the genes having closely related functions with patient’s phenotype and also not extensively covered by CNVs in Database of Genome Variants. We performed Sanger sequencing for these seven variants in patient’s family (the father and son having the same translocation and the father also having dyslexia and stutter). Four variants were found to be false positive. Both the patient and healthy mother possessed two variants. All three family members possessed one variant. Overall, we have not found a promising disease causing variants other than the translocation found in patient CMS12200.

#### Histone H3 RIP-qRT-PCR (related to Figure 2F)

Approximately 20-50×10^6^ cells were used for each experiment. Cells were crosslinked with 1% formaldehyde. Cell pellet was resuspended in equivalent volume of Nuclear Lysis Buffer (i.e. 100mg-1mL buffer) (50 mM Tris-Cl pH 7.0, 10 mM EDTA, 1% SDS, 100x PMSF, 50x protease inhibitors, and 200x Superase inhibitor). Chromatin was sheared using Covaris sonicator until DNA was fragmented to 200-1000 bp range and diluted 2-fold using Dilution Buffer (0.01% SDS, 1.1% Triton X 100, 1.2 mM EDTA, 16.7 mM Tris-Cl pH 7.0, 167 mM NaCl, 100x PMSF, 50x protease inhibitors, and 200x Superase inhibitor). Samples were incubated with 5 µg of H3 or IgG antibody rotating overnight at 4°C. Protein A dynabeads (50uL) were washed in Dilution buffer and added to the chromatin for 2 hours rotating at room temperature. Immunoprecipitate fraction was washed four times with Wash buffer (100 mM Tris-Cl pH 7.0, 500mM LiCl, 1% NP40, 1% sodium deoxycholate, and PMSF). Subsequently, the immunoprecipitate fraction was eluted from beads by vortexing for 30 minutes at room temperature using elution buffer (50mM sodium bicarbonate, and 1% SDS). Immediately after 5% of 3M Sodium Acetate was added to neutralize pH. Proteinase K treatment proceeded for 45 minutes at 45C, followed by RNA extraction using Trizol. Isolated RNA was subjected to DNAse treatment using TurboDNase and purified by phenol-chloroform extraction and ethanol precipitation. For qRT-PCR analysis we used Roche’s Lightcycler and Stratagene’s RT kit.

#### RNA-seq library preparation (related to Figure 3E, 3G, S7C)

We followed protocols previously described (Wapinski et al., 2013). Briefly, for the RNA-sequencing experiment in figure 3, 4 and S7, libraries were produced from poly-A enriched mRNA using TruSeq kit (Illumina). They were subsequently sequenced using the NextSeq or HiSeq platform producing paired ends reads.

#### RNA-seq analysis for loss and gain of function analysis (related to Figure 3E, 3G, S7C)

Reads obtained were first mapped using Tophat. Expression for each gene was calculated using Cuffdiff (Figure 3E, Figure 3G) or DEGSeq (Figure S7C) using default settings. For DEGSeq briefly, only properly paired mapped reads were used ^60^. DEGSeq selected longest transcript for each gene, when multiple isoforms were found. Raw counts for each sample were merged into a table and transformed to logarithmic scale. Batch effect among samples was removed using ComBat method in sva package in R ^61^. Subsequently, expression values were transformed raw counts and differentially expressed genes were identified by DESeq2 package by comparing different conditions using default parameters ^61^. Gene ontology analyses were performed using PANTHER/DAVID.

#### ChIRP-seq and data analysis (related to Figure 4)

To determine the genome-wide localization of *lnc-Nr2f1*1 we followed protocols previously described (Chu et al., 2011)^51^. ChIRP was performed using biotinylated probes designed against mouse lnc-Nr2f1 using the ChIRP probes designer (Biosearch Technologies). Independent even and odd probe pools were used to ensure lncRNA-specific retrieval (Refer to separate document for odd and even sequences targeting human and mouse lnc-Nr2F1). Mouse ES-iN samples are crosslinked in 3% formaldehyde. RNase pre-treated samples are served as negative controls for probe-DNA hybridization. ChIRP libraries are constructed using the NEBNext DNA library preparation kit (New England Biolabs). Sequencing libraries were barcoded using TruSeq adapters and sequenced on HiSeq or NextSeq instruments (Illumina). Reads were processed using the ChIRP-seq pipeline^59^. Even-odd ChIRP-seq tracks are merged as previously described ^59^. Peaks were called from the merged tracks over RNase control tracks using MACS14. Overlapping peaks from all replicates were final peaks. High confidence peaks were then filtered by their significance [-log10 (p-value) ≥100] and correlation between even/odd probes >0, average coverage (>2 for mES-iN, >1 for hNPC). For hNPC ChIRP of long and short isoforms, the analysis pipeline and filtering criteria are the same. Sequence motifs were discovered using Homer in 200-bp windows. Peak associated gene sets were obtained through GREAT ^52^ (http://great.stanford.edu/). Peaks are assigned to genes according to whether peaks are in gene’s regulatory domain. Gene regulatory domain is defined as: Each gene is assigned a basal regulatory domain of a distance 5kb upstream and 1kb downstream of the TSS (regardless of other nearby genes). The gene regulatory domain is extended in both directions to the nearest gene’s basal domain but no more than the 1000kb extension in one direction. Gene Ontology of gene sets were performed using Metascape (http://metascape.org/). For overlapping mouse ChIRP-seq peak with chromatin features (CTCF, enhancer, H3K27ac, H3K4me3 and PolII) in mouse cortex and E14.5 brain, chromatin annotation files are obtained from public available Chip-seq data from Bing Ren lab ^51^. For overlapping human ChIRP-seq peak with chromatin features, ChromHMM model of 25 chromatin states and 12 histone modification marks in neuron cells was used^62^. In addition, peaks are annotated according to distance to genes in figure 6 (promoter: −2kb to +1kb of TSS, enhancer: −2kb to −10kb of TSS, exon: exon of a gene, intron: intro of a gene, gene tail: 0 to 2kb downstream of the end of a gene, intergenic: none of the above).

### Biochemistry

#### Single molecule RNA FISH protocol and probes

Probes were designed using Stellaris probe designer tool and synthesized by Stellaris. Adherent cells were grown in 12mm coverglass, fixed in 1% formaldehyde for 10 minutes at room temperature, washed twice with phosphate buffer saline (PBS), and permeabilized using 70% ethanol at 4C overnight. Fixed cells were subjected to RNAse treatment for 30 minutes at 37C with 0.1mg/mL RNAse A. After washing (2x SSC, 10% Formamide), hybridization with 250nM probes in hybridization buffer (10% dextran sulfate, 10% formamide, 2x SSC) at 37C overnight in a coverglass protected from light. The next day, washed (2x SSC, 10% Formamide) at 37C for 30 minutes. DAPI staining was added to a clean coverslip and coverglass mounted. Slides were images using confocal microscopy.

#### In situ hybridization

E13.5 mouse embryos were fixed at 4°C with 4% (weight/volume) paraformaldehyde in PBS overnight. Samples were cryoprotected overnight with 30% (weight/volume) sucrose in PBS, embedded in OCT (Tissue-Tek), and frozen on dry ice. Frozen embryos were sectioned on a cryostat at 16μm. Sections were processed for in situ hybridization. Frozen sections were treated sequentially with 0.3% (volume/volume) Triton-X in PBS and RIPA buffer (150mM NaCl, 50mM Tris-HCl (pH 8.1), 1mM EDTA, 1% NP-40, 0.5% Sodium Deoxycholate, 0.1% SDS). Sections were postfixed in 4% paraformaldehyde at room temperature for 15 minutes and washed with PBS. Subsequently, the sections were treated with 0.25% acetic anhydride in 0.1M triethanolamine for 15 minutes and washed with PBS. Sections were incubated in hybridization buffer (50% formamide deionized, 5× SSC, 5× Denhardts, 500μg/mL Salmon Sperm DNA, 250μg/mL yeast tRNA) containing DIG-labeled probes at 65°C overnight. Hybridized sections were washed two times in washing solution (2× SSC, 50% formamide, 0.1% Tween 20) at 65°C for 60 minutes. After washing, sections were incubated for 1 hour in 1% (weight/volume) blocking reagent (0.1M Maleic Acid, 0.15M NaCl (pH 7.5), 0.1% Tween 20, Roche). Subsequently, incubated with an alkaline phosphatase (AP)-coupled antibody (Roche) at 4°C overnight. After rinsing, the signals were visualized with nitro-blue tetrazolium chloride (NBT)/5-Bromo-4-Chloro-3′-In-dolylphosphatase p-Toluidine salt (BCIP) (Sigma). The DIG-labeled antisense RNA probe for detecting mouse NR2F1 corresponds to the CDS region and for *lnc-Nr2f1* corresponds to 800bp region. The DIG-labeled sense RNA probe for both, NR2F1 and *lnc-Nr2f1*, corresponds to the same region as the antisense probe in the reverse direction. Probes were generated by in vitro transcription with T7 RNA polymerase (Roche) using the DNA templates containing a promoter sequence of T7 RNA polymerase promoter (TAA TAC GAC TCA CTA TAG GG) followed by a complimentary sequence of target RNA. DNA templates were amplified by PCR with the following primers: For *lnc-Nr2f1* probe: (F) GTG GCC ATG GAA TGG TGT AGC AGA, and (R) GTC TGA GTG TTT GTT TGA CTG AAT GT; NR2F1 probe: (F) CGG TTC AGC GAG GAA GAA TGC CT, and (R) CTA GGA ACA CTG GAT GGA CAT GTA AG.

#### Cellular fractionation

Cell fractionation of primary neocortical cells (prepared from E12.5 mouse cerebral cortex) into cytoplasmic and nuclear RNA fractions was performed with a nuclear/cytoplasm fractionation kit (PARIS kit, Ambion) following the instructions of the manufacturer. The amount of RNA in each fraction was determined by qRT-PCR in a Roche LightCycler with Brilliant III Ultra-Fast SYBR Green QRT-PCR Master Mix (Agilent). For primer sequences refer to separate document.

#### Immunofluorescence

Cells were fixed with 4% paraformaldehyde for 15 minutes and subsequently lysed and blocked with blocking buffer [PBS + 0.1% Triton X (Sigma Aldrich) + 5% Cosmic calf serum (Thermo Scientific)] for 30 minutes. Primary antibodies diluted with blocking buffer were added to the wells and left for an hour. The following antibodies were used for immunostaining: mouse anti-MAP2 (Sigma, 1:500), rabbit anti-Tuj1 (Covance, 1:1000), goat anti-Sox1 (R&D, 1:100) and mouse anti-hNestin (R&D, 1:1000). The wells were subsequently washed three times with the blocking buffer. Secondary antibodies conjugated with Alexa dyes (1:1000, Invitrogen) diluted with the blocking buffer were added to the wells and left for an hour. The wells were again washed three times with the blocking buffer. 4’,6-Diamidino-2-phenylindole (DAPI) (Life Technologies, 1: 10,000) diluted in PBS was added for 1 minute for nuclear staining.

#### Western blotting

Cells were lysed with 1 volume of RIPA buffer with cOmplete protease inhibitors (Sigma-Aldrich) and equivolume of 2x Laemmli buffer was added. The samples were then boiled for 5 minutes at 95^°^C and subsequently separated in 4-12% Bis-Tris gel with MES buffer (Invitrogen) and transferred onto PVDF membrane for 2 hours at 4^°^C. Blots were then blocked in blocking buffer (PBS + 0.1% Tween-20 (Sigma-Aldrich) + 5% fat-free milk) for 30 minutes and subsequently incubated overnight with primary antibodies at 4^°^C. The primary antibodies used are rabbit anti-HSP90 (Cell Signalling) and rabbit anti-NR2F1 (Cell Signalling). The blots were washed three times in PBS + 0.1% Tween-20 for 10 minutes each. Next, the blots were incubated with secondary antibodies conjugated to horseradish peroxide (Jackson immunoresearch) were diluted in blocking buffer for 1 hour. The blots were washed three times in PBS + 0.1% Tween-20 for 10 minutes each and once with PBS before adding chemiluminescence substrates (Perkin Elmer) for signal detection on films.

---

**Fig. S1.**
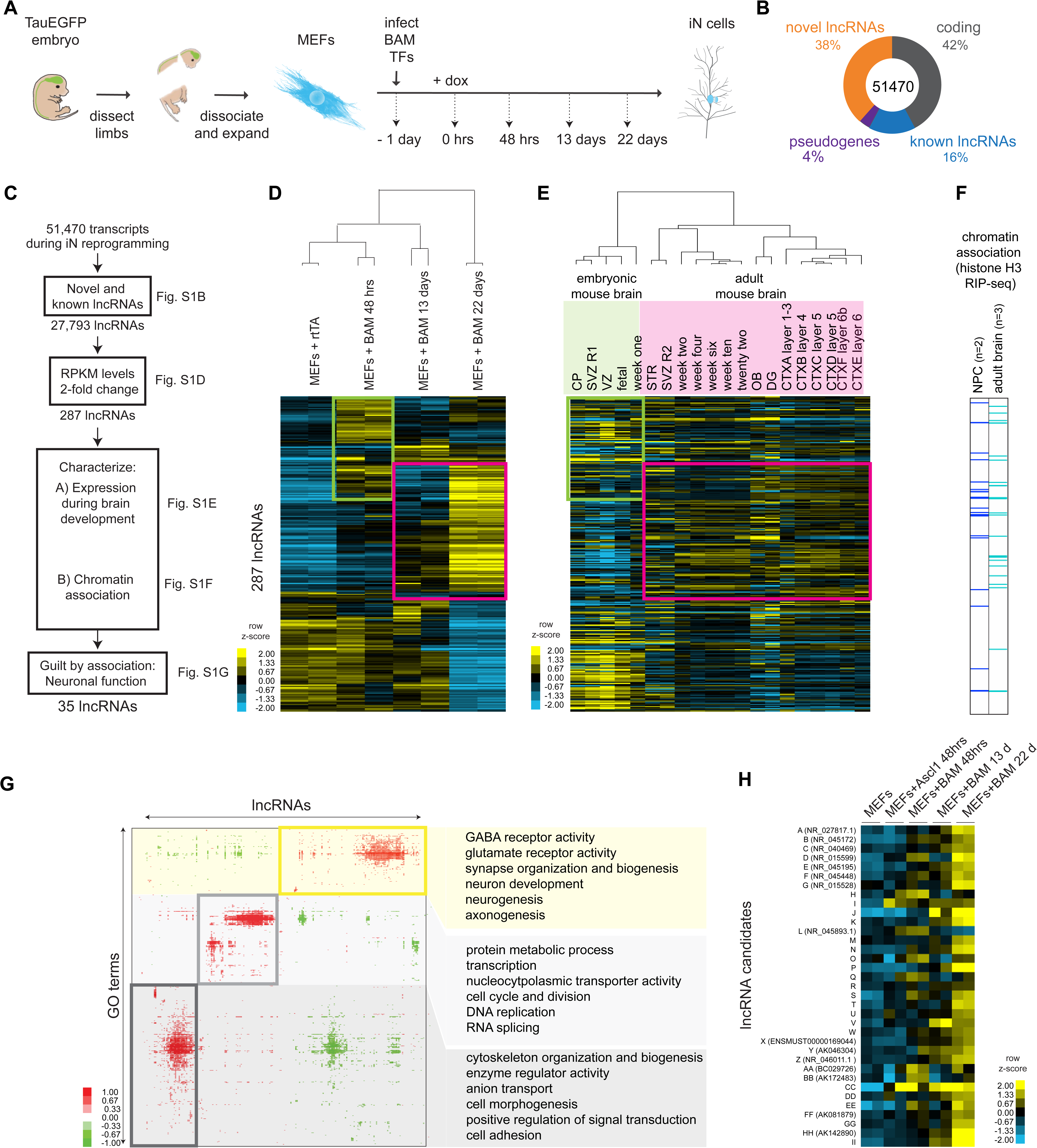
Molecular profiling of direct fibroblast to iN cell reprogramming nominates functional lncRNAs involved in neurogenesis. (A) Schematic representation of experimental time points generated for this study. (B) Classification of the iN cell transcriptome consisting of 51,470 transcripts based on coding genes and non-coding RNAs. (C) Diagram depicting pipeline derived in this study to enrich for candidate lncRNAs with strong neuronal association. (D) Hierarchical clustering heatmap of lncRNA expression during iN cell reprogramming by RNA-seq across indicated time points (n=2 biological replicates). Shown are 287 lncRNAs that changed expression at least two-fold at any time point (p<0.05). Fold change is represented in logarithmic scale normalized to the mean expression value of a gene across all samples. The green box highlights the genes which are upregulated in the MEF + BAM 48 hours. The same set of genes are upregulated in embryonic mouse brain (see Figure S1E). The pink box highlights the genes which are upregulated in the MEF + BAM 22 hours. The same set of genes are upregulated in adult mouse brain (see Figure S1E). (E) Hierarchical clustering heatmap of iN lncRNAs from Figure S1D across mouse brain tissues from publicly available data. Expression levels are represented in logarithmic scale normalized to the mean expression value of a gene across all samples. (F) Chromatin association of iN lncRNAs determined by histone H3-RIP-seq in Neuronal Progenitor Cells (NPCs) and total adult brain (n=2 biological replicates for NPCs and n=3 for mouse brain) presented in binary format. Shown are lncRNAs that have significant enrichment over background (>2-fold, p<0.05) and consistent enriched in the chromatin among biological replicates. (G) Co-expression analysis using Genomica between iN lncRNAs and mRNAs associated with Gene ontology (GO) terms. Highlighted in the yellow box are lncRNA candidates which are associated with neuronal GO terms. (H) RNA-seq heatmap of 35 filtered candidate lncRNAs across MEF-to-iN cell reprogramming. In brackets are the Refseq ID for the annotated lncRNAs. See the supplementary documents for the sequences and coordinates of the 35 candidates.

**Fig. S2.**
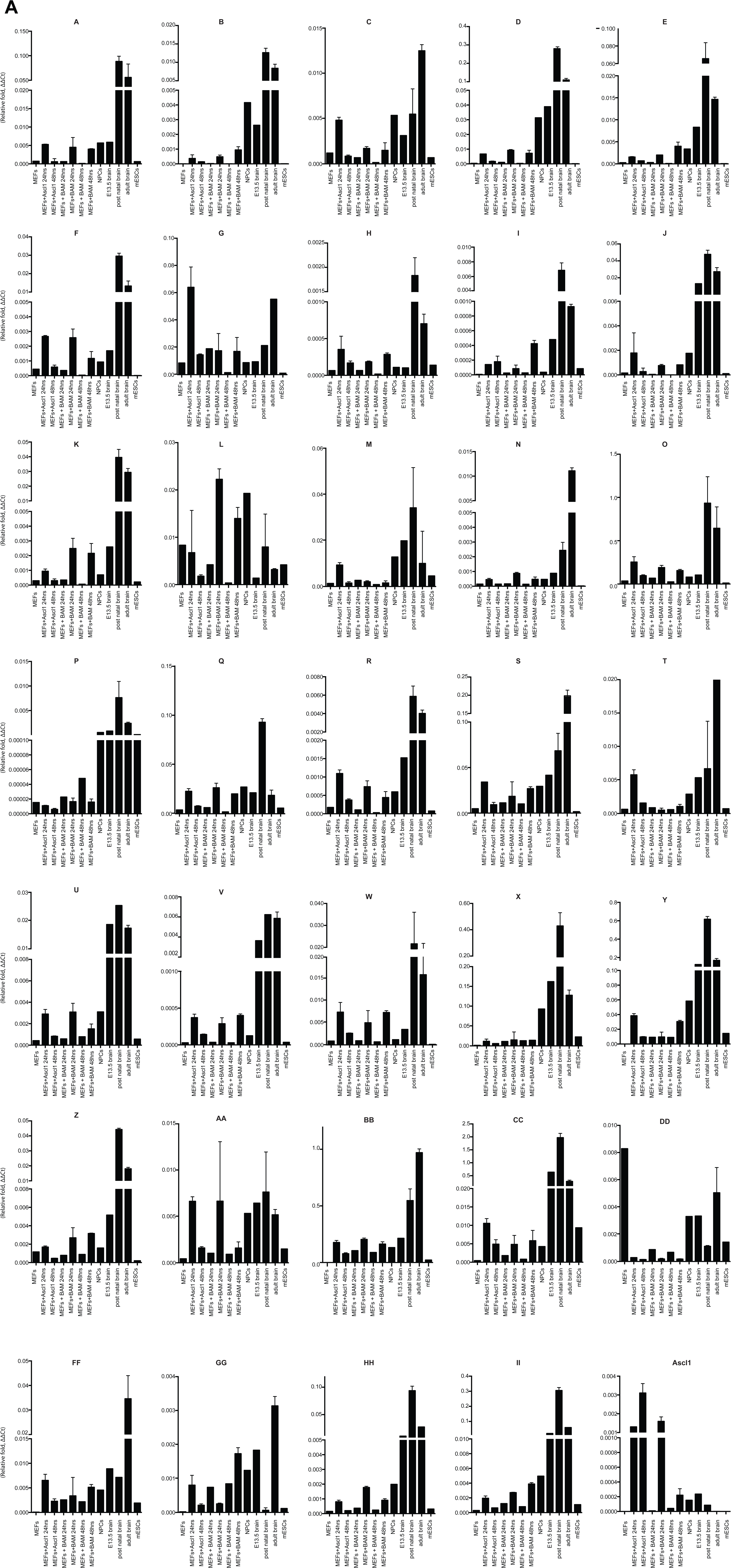
QRT-PCR validation of candidate lncRNAs expression, Related to Figure 1. (A) Expression detection of candidate lncRNAs by qRT-PCR across early stages of iN cell reprogramming and mouse brain development.

**Fig. S3.**
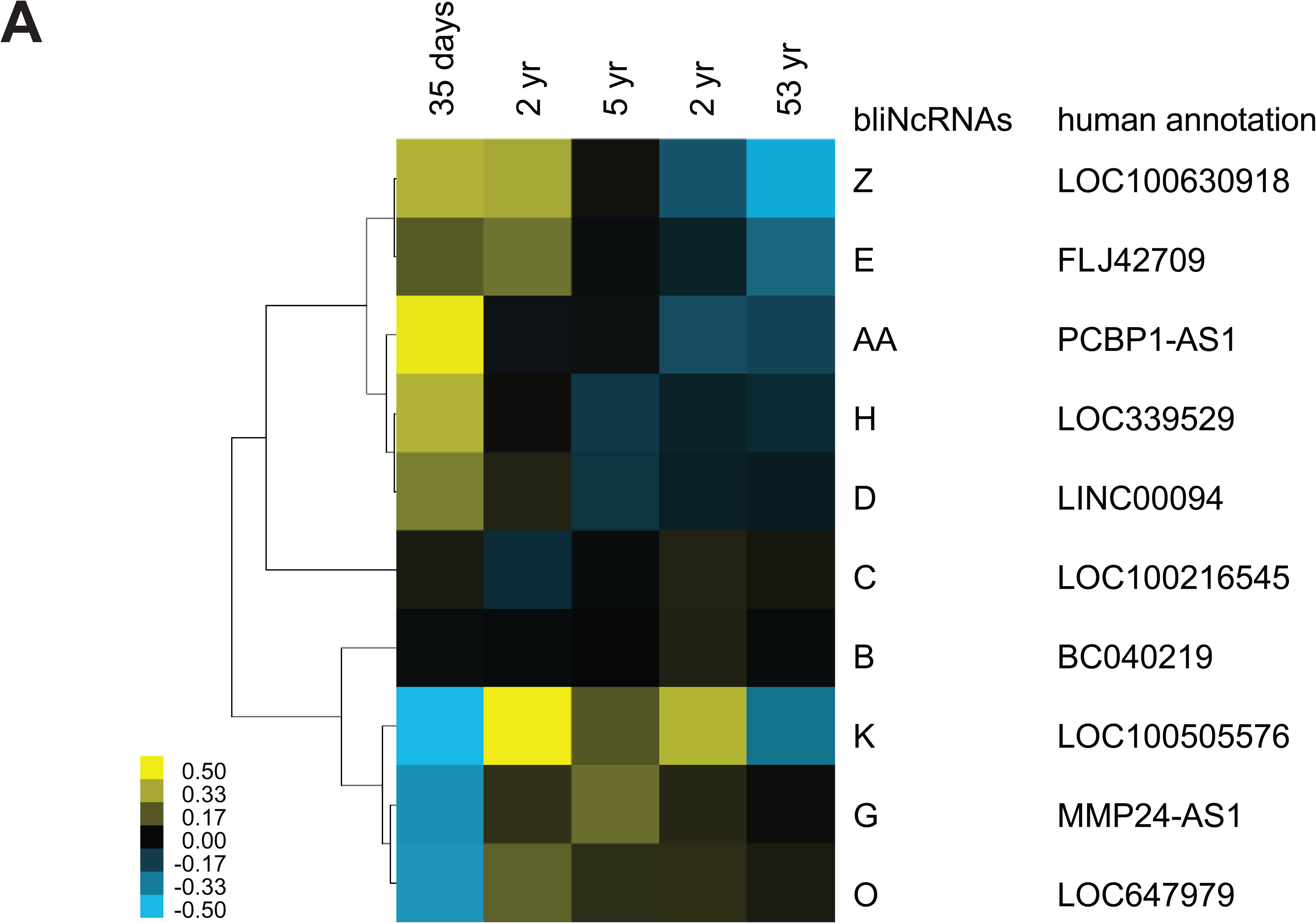
Conserved lncRNAs have distinct pattern of expression across different stages of the human brain, Related to Fig. 1. (A) Heatmap of annotated syntenic lncRNAs across human brain development detected by RNA-seq. Expression is represented in logarithmic scale normalized to the mean RPKM value of a gene across all samples. See Supplementary documents for the lncRNA sequences.

**Fig. S4.**
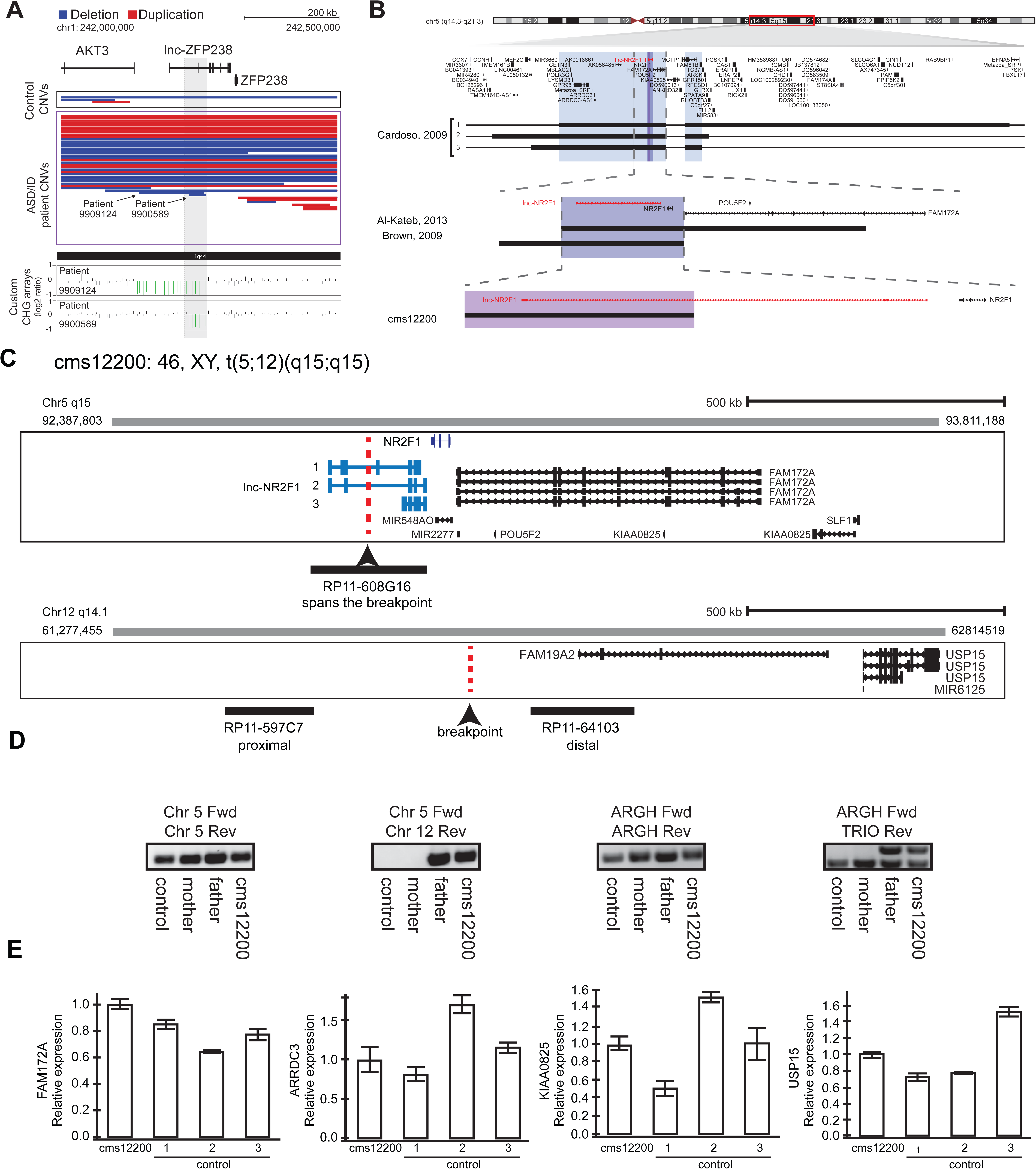
Other reports of CNVs affecting *lnc-NR2F1* and an example of focal deletion affecting *lnc-*ZFP238 and characterization fo patient CMS12200, Related to Fig. 1. (A) Top: Representative tracks for lncRNA H locus, also known as *lnc-*ZFP238. Depicted in blue are deletions and in red duplications. Arrow points to two significant focal deletions in two distinct patients. Bottom: Custom CGH arrays used to validate chromosomal aberration in patients 9909124 and 9900584 harboring focal deletions represented in green signal. (B) Representative tracks of previously reported deletions affecting chromosome region 5q15 in patients with neurodevelopmental and neuropsychiatric disorders. The black rectangle represents affected genomic region corresponding to the patient described in a previous publication to the left. Top panel: light blue box represents minimal common region amongst three patients in Cardoso et al, 2009 report. There are multiple genes affected in the locus. Middle panel: purple box depicts common deleted region reported amongst two patients from Brown, et al 2009, and Al-Kateb, et al 2013 studies. The region encompasses only two genes: *NR2F1* and *lnc-NR2F1*. Bottom panel: Patient CMS 12200 described in this study harboring a balanced chromosomal translocation disrupting *lnc-NR2F1* only as depicted by the pink box. In red, *lnc-NR2F1* is highlighted. (C) Schematic representation of breakpoint region between chromosome 5 and 12 for patient CMS12200. Illustrated is also the probe design to confirm event. Probe RP11-608G16 spans the breakpoint in chromosome 5. Probe RP11-597C7 is proximal to the breakpoint on chromosome 12. Probe RP11-64103 is distal to the breakpoint on chromosome 12. (D) Genomic PCR to confirm t(5;12) translocation using primers spanning control region in chromosome 5 (left), translocation between chromosome 5 and 12 (middle left), unaffected region in ARGH gene (middle right), and non-pathological gene duplication for TRIO exon in ARGH intronic loci (right). Samples from control (GM12878), mother (does not harbor translocation), father (t(5;12)), and CMS12200 patient (t(5;12)). (E) Expression of genes proximal to breakpoint is unaffected as measured by RT-qPCR in CMS12200 patient lymphocytes and three distinct control samples.

**Fig. S5.**
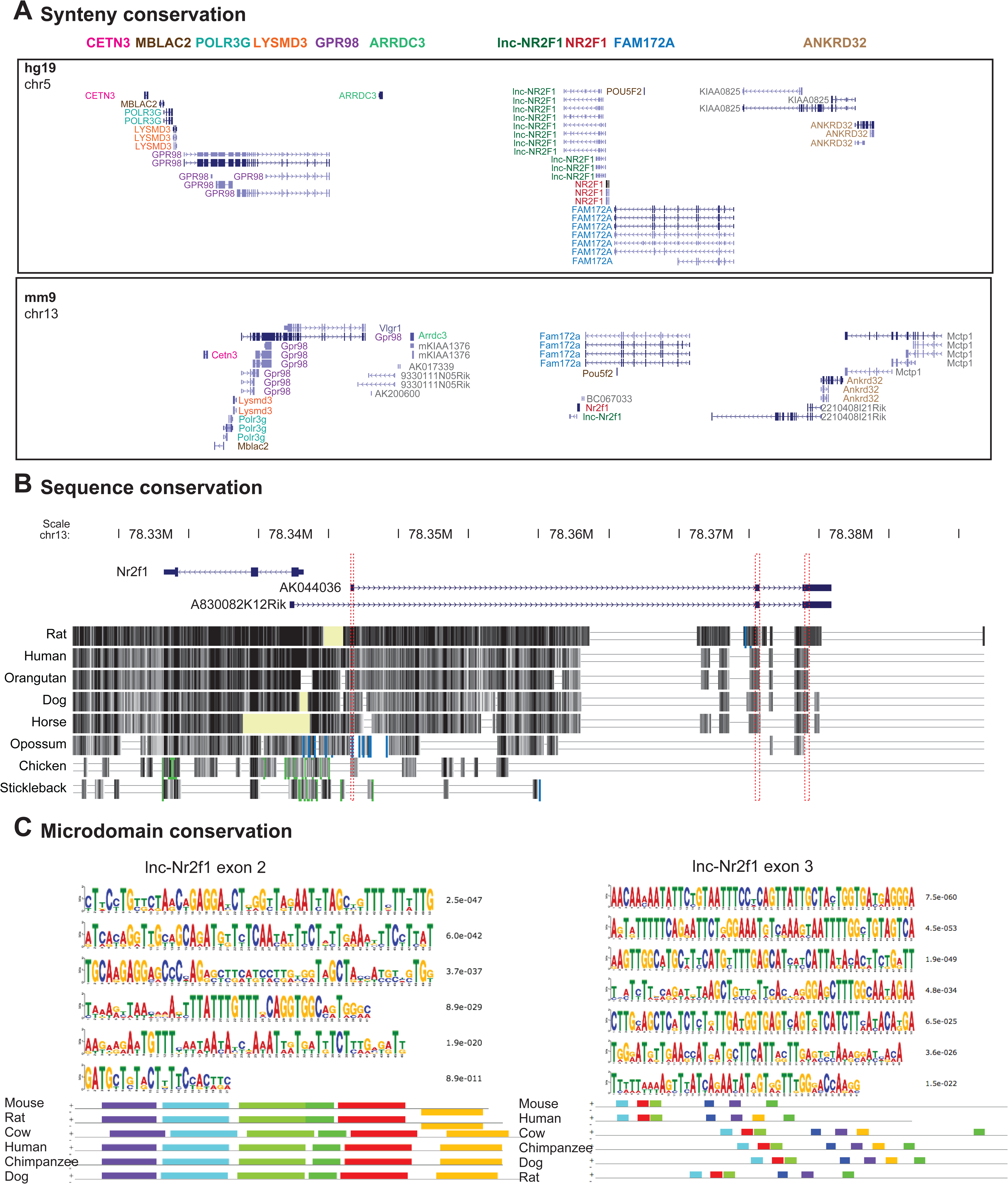
Synteny, sequence and microdomain conservation of lnc-*NR2F1*, Related to Fig. 2. (A) UCSC browser track for human (top) and mouse (bottom) showing synteny conservation around the lnc-*Nr2f1 – Nr2f1* locus. The same gene between the two species is color-coded (B) UCSC browser track showing the sequence conservation of the three exons (highlighted in red) across different species. (C) Sequences around the conserved exons show short sequence homology from different species. MEME (http://meme-suite.org/tools/meme) is used to discover motif homology. See Supplementary documents for the sequences used.

**Fig. S6.**
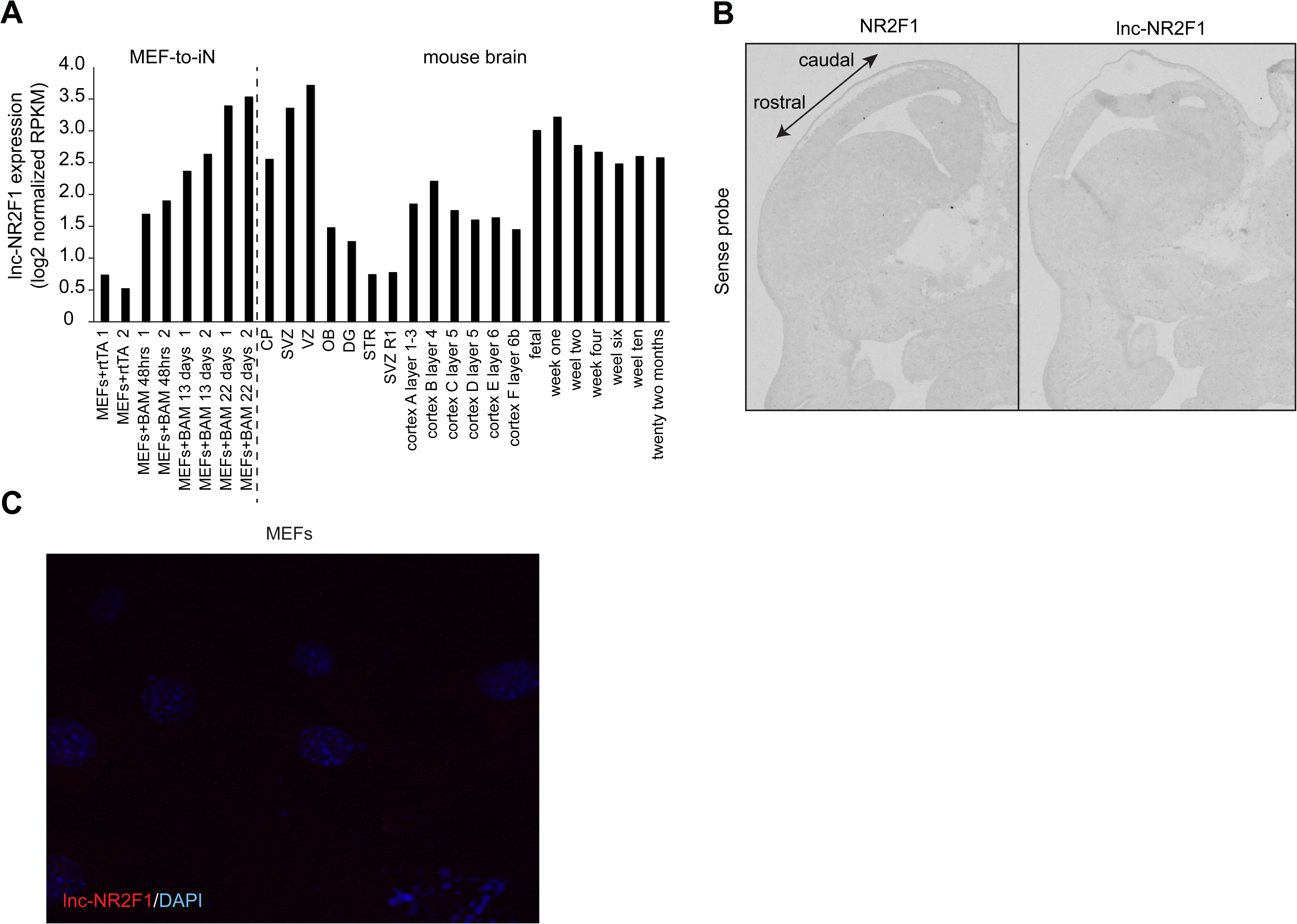
Characterization of *lnc-Nr2f1* localization, Related to Fig. 2. (A) RPKM counts for *lnc-Nr2f1* during MEF-to-iN-reprogramming and across different stages and tissues of mouse brain development. (B) Control in situ hybridization for NR2F1 and *lnc-Nr2f1* using sense probe. (C) Control single molecule RNA-FISH in MEFs infected with rtTA alone in order to determine background signal.

**Fig. S7.**
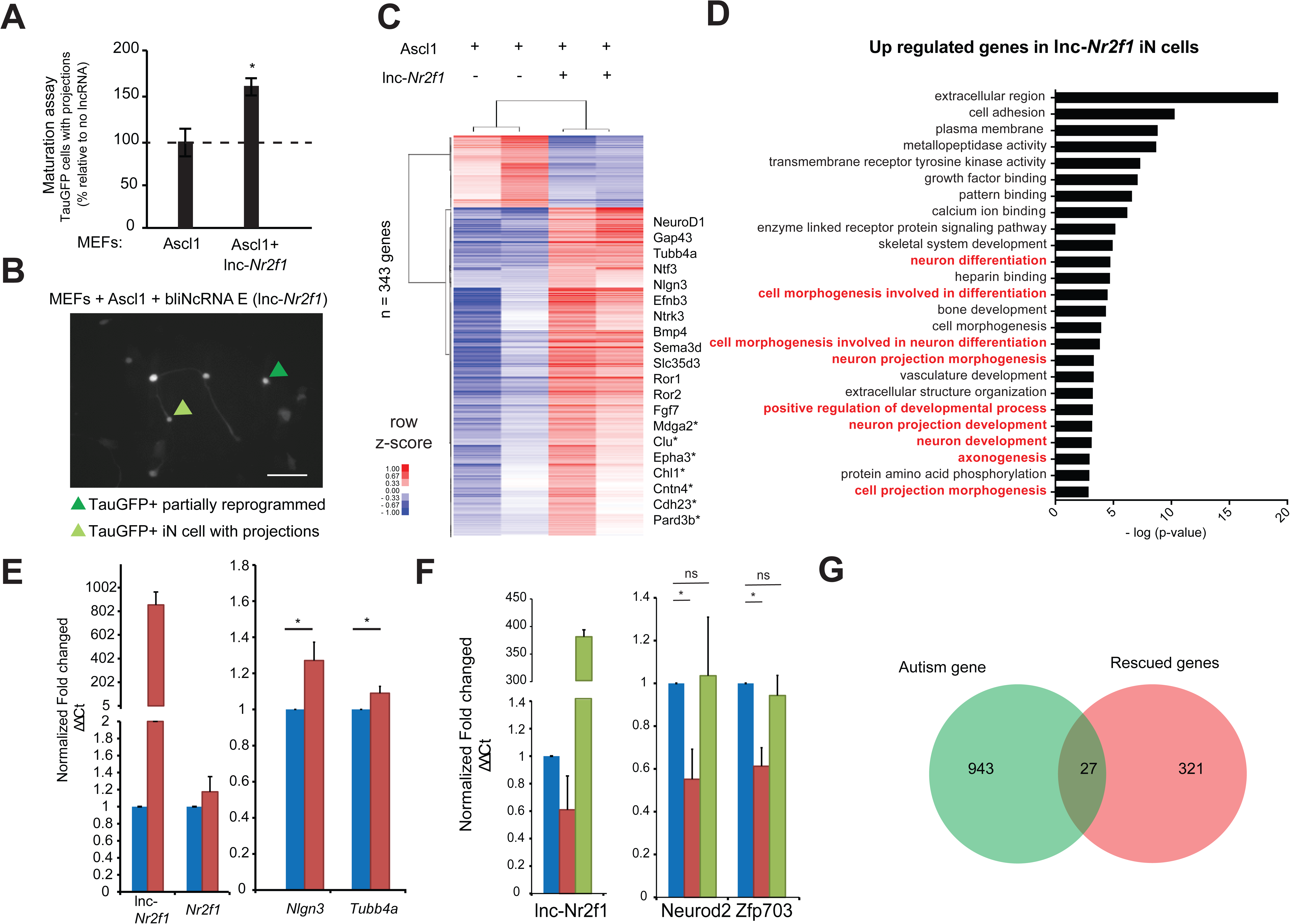
Characterization of the roles of *lnc-Nr2f1* during iN reprogramming, Related to Fig. 3. (A) Percentage of TauGFP positive cells with projections normalized to number of TauGFP cells. TauGFP cells with projections longer than three times the diameter of the cell body were counted and normalized to the total number of TauGFP positive cells. The sequence for mouse *lnc-Nr2f1* is available in the supplementary documents (n=4, Student t-test, Two tailed, Error bars show s.e.m). (B) Immunofluorescence staining depicting how a TauGFP positive cell with projections (example highlighted in light green) is differentiated from a TauGFP positive partially reprogrammed iN cell (example highlighted in dark green) at 7 days. Scale bar = 50µm. (C) Hierarchical clustering heatmap of differentially expressed genes detected by RNA-seq in MEFs expressing Ascl1 alone compared to Ascl1 and *lnc-Nr2f1* after 7 days (n=2 biological replicates, FDR corrected p<0.001). Shown are 343 genes. 311 genes are upregulated and 32 genes are downregulated. Fold change is represented in logarithmic scale normalized to the mean expression value of a gene across all samples. Representative gene names are included. Those with (*) have been linked to neurological disorders curated by Basu et al. (D) Gene ontology of the upregulated genes upon *lnc-Nr2f1* overexpression in Ascl1 MEF-iN 7 days. Highlighted in red are neuronal GO terms. (E) qRT-PCR validation downstream neuronal genes of the RNA-sequencing results in Fig. S3C. Ectopic expression of *lnc-Nr2f1* led to upregulation of several downstream neuronal genes. (n=3, * indicates p<0.05). Error bars show s.e.m (F) qRT-PCR validation of several target genes in Fig. 3G that go down when *lnc-Nr2f1* is knocked out which can be subsequently rescued with *lnc-Nr2f1*. (n=4, * indicates p<0.05). Error bars show s.e.m. (G) Venn diagram representing the overlap between autism related genes and rescued genes from figure 4g.

**Fig. S8.**
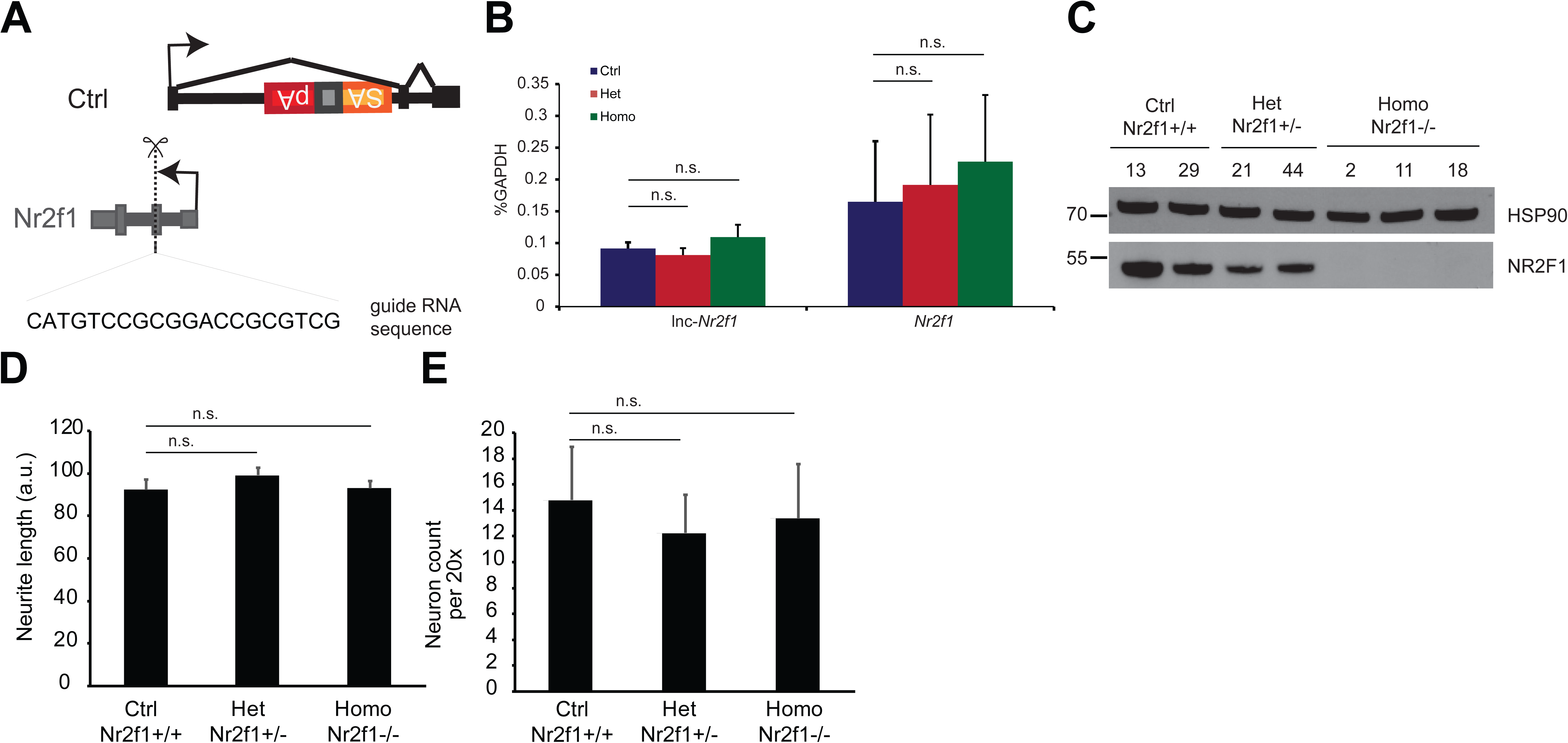
Characterization of the epistasis relationship between mouse *Nr2f1* and *lnc-Nr2f1*, Related to Fig. 3. (A) CRISPR knock out strategy to generate *Nr2f1* knockout (Homo) and heterozygous null lines (Het) from the control mES cells (Ctrl). (B) qRT-PCR results for *lnc-Nr2f1* and *Nr2f1* in the Ctrl, *Nr2f1* heterozygous null and *Nr2f1* knock out day 4 Ngn2 mES-iN. (n=4 for Ctrl and Homo, n=6 for Het; n.s. denotes not significant by two tailed t test) (C) Western blot showing the level of NR2F1 for individual clones of Ctrl, Het and Homo for day 4 Ngn2 mES-iN. (D) Neurite length measurement of the Ngn2 day 3 mES iN cells generated from the *Nr2f1* Ctrl, Het or Homo lines. (n=4 for Ctrl and Homo, n=6 for Het) (n.s. indicates p<0.05). (E) Number of neurons per 20x the Ngn2 day 3 mES iN cells generated from the *Nr2f1* Ctrl, Het or Homo lines. (n=4 for Ctrl and Homo, n=6 for Het) (n.s. indicates p<0.05).

**Fig. S9.**
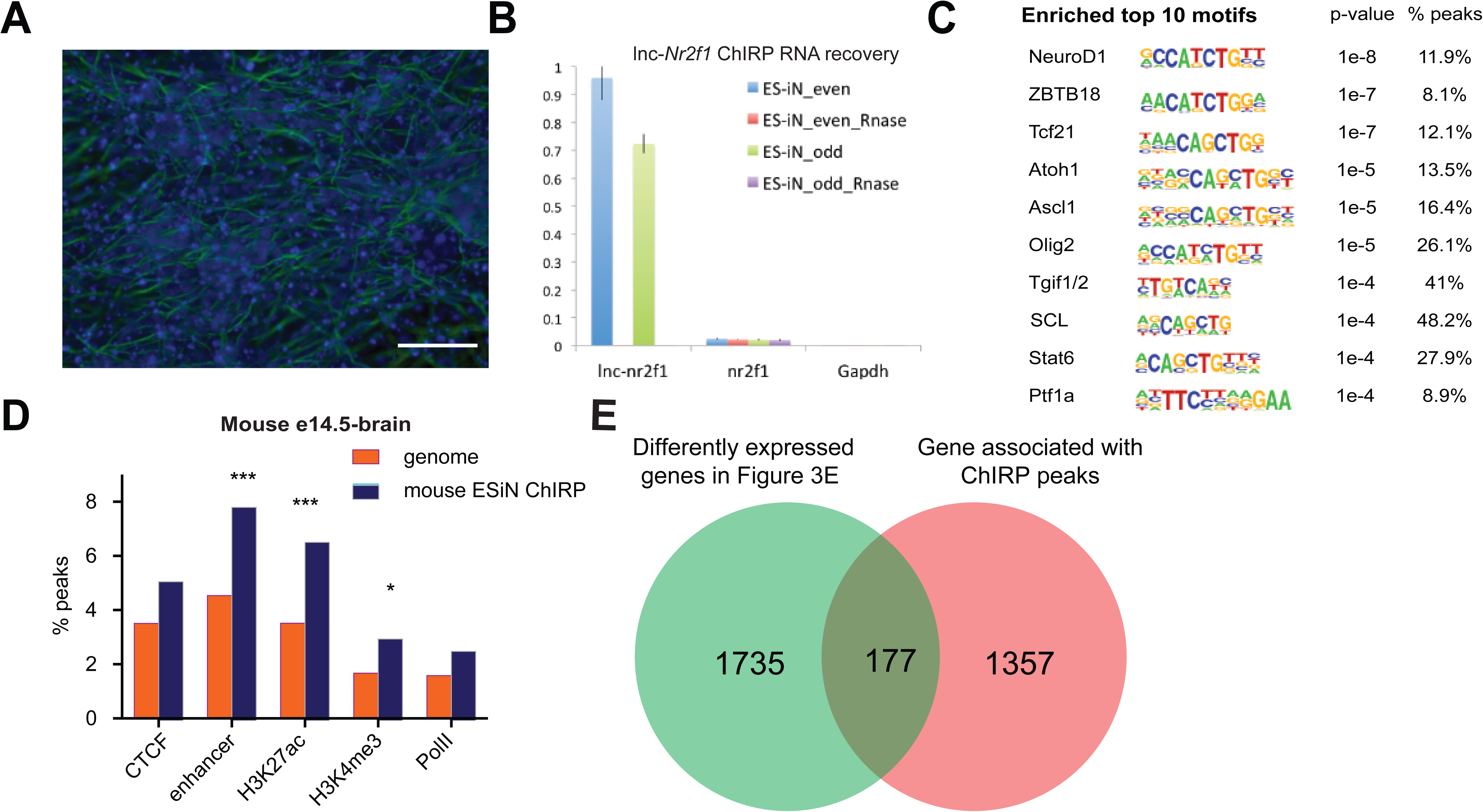
Identification of *lnc-NR2F1* role in transcriptional regulation, Related to Fig. 4. (A) Immunofluorescence staining for the day 4 mouse embryonic stem cell derived induced neurons (mES-iN) used in the *lnc-Nr2f1* ChIRP. (Green=β-III-tubulin, Blue=DAPI) (Scale bar=50µm) (B) *Lnc-Nr2f1* RNA pull down efficiency for both even and odd probes (C) The enriched motifs with their corresponding p-value and the percentage of peaks with the given motifs. (D) Percentage of mES-iN ChIRP-seq peaks which overlap with CTCF, enhancer, H3K27ac, H3K4Me3 and PolII defined in mouse E14.5 brain relative to the control. (*** represents p<0.0001, * represents p<0.05, Chi-square test) (E) Venn diagram showing the overlap between 1912 genes from Figure 4E and the 1534 genes adjacent to the 1092 high confident mES-iN ChIRP peaks. (p<0.0001, Chi square test)

**Fig. S10.**
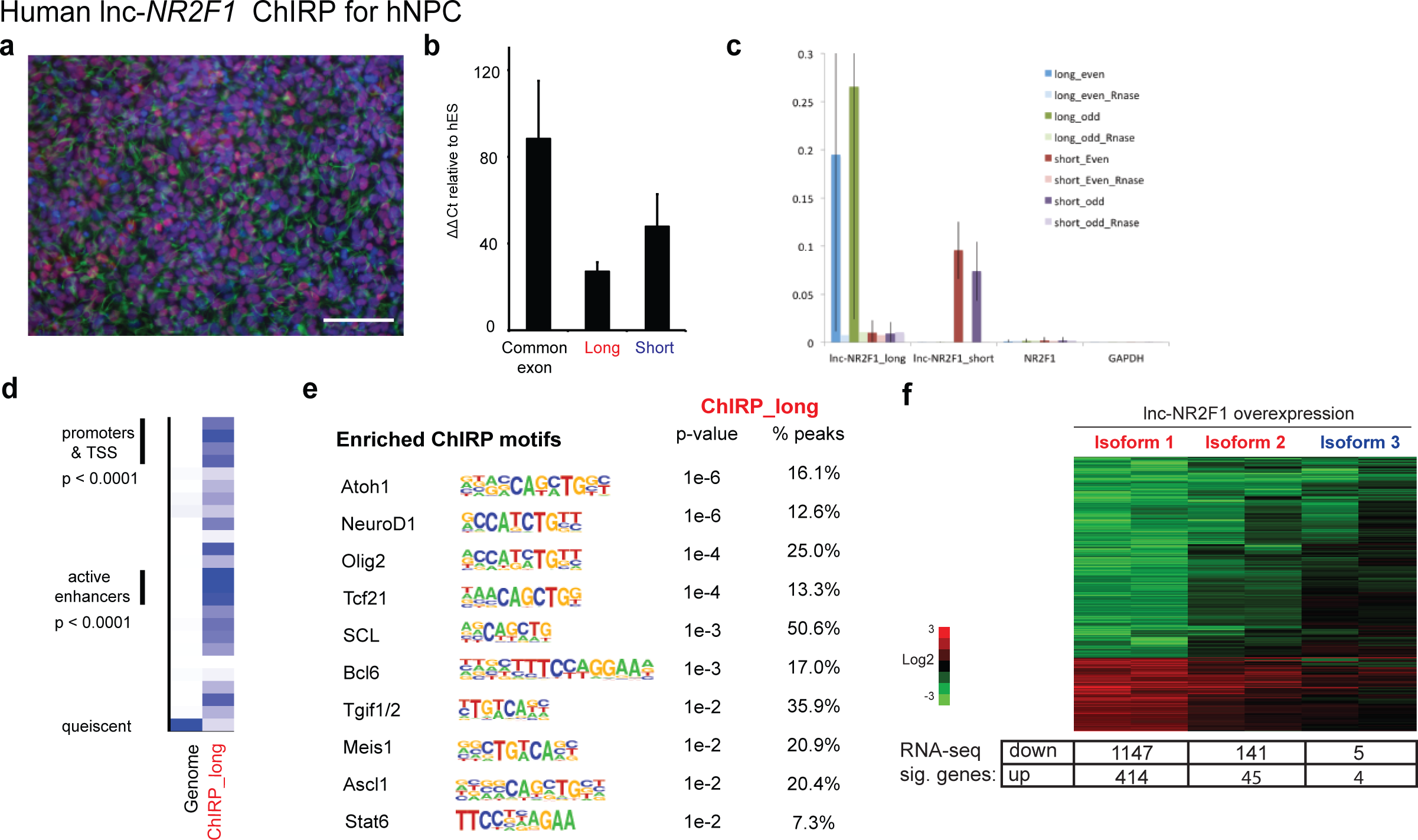
Identification of *lnc-NR2F1* role in transcriptional regulation, Related to Figure 5. (A) Immunofluorescence staining for day 12 human neural progenitor cells (hNPC) differentiated from HUES9 cells using dual SMAD protocol. (Green=human NESTIN, Red=human SOX1, Blue=DAPI) (Scale bar=50µm) (B) qRT-PCT using primers specific to the common, long or short exon in day 12 hNPC. (n=3). (C) *Lnc-NR2F1* RNA pull down efficiency for the long and short isoform-specific exons of for both even and odd probes. The pull down is specific to *lnc-NR2F1* since there is very little *NR2F1* or *GAPDH*. (D) ChromHMM model ran on the peaks from the high confident long isoform-specific exon (exon 11) ChIRP peaks performed in hNPC. The peaks are enriched in the promoters and TSSs, active enhancers and quiescent chromatin regions. (E) The enriched transcription factor motifs for the long isoform-specific domain ChIRP experiments in hNPC. (F) Heatmap showing the gene expression changes in a human neuroblastoma cell line (SK-N-SH) upon overexpression of human *lnc-NR2F1* isoform 1, 2 and 3 respectively normalized to the control (>2-fold, p<0.05, FDR<0.05). The box below the heatmap shows the number of genes significantly upregulated or downregulated.

**Table S1. Diagnostic comparison between studies of patients with affected *lnc-NR2F1* locus. Related to Fig. 2 (See Supplementary Document)** (A) Summary of diagnosis for previously reported patients, including patient CMS12200 described in this study. Highlighted in grey are the shared diagnostic features across patients. Adapted figure ^39^.

## References

1. Flynn, R.A. & Chang, H.Y. Long noncoding RNAs in cell-fate programming and reprogramming. Cell Stem Cell 14, 752–61 (2014).

2. Wapinski, O. & Chang, H.Y. Long noncoding RNAs and human disease. Trends Cell Biol 21, 354–61 (2011).

3. Fertuzinhos, S. et al. Laminar and temporal expression dynamics of coding and noncoding RNAs in the mouse neocortex. Cell Rep 6, 938–50 (2014).

4. Valadkhan, S. & Nilsen, T.W. Reprogramming of the non-coding transcriptome during brain development. J Biol 9, 5 (2010).

5. Lv, J. et al. Long non-coding RNA identification over mouse brain development by integrative modeling of chromatin and genomic features. Nucleic Acids Res 41, 10044–61 (2013).

6. Aprea, J. et al. Transcriptome sequencing during mouse brain development identifies long non-coding RNAs functionally involved in neurogenic commitment. EMBO J 32, 3145–60 (2013).

7. Ramos, A.D. et al. Integration of genome-wide approaches identifies lncRNAs of adult neural stem cells and their progeny in vivo. Cell Stem Cell 12, 616–28 (2013).

8. Ramos, A.D. et al. The long noncoding RNA pnky regulates neuronal differentiation of embryonic and postnatal neural stem cells. Cell Stem Cell 16, 439–47 (2015).

9. Ng, S.Y., Bogu, G.K., Soh, B.S. & Stanton, L.W. The long noncoding RNA RMST interacts with SOX2 to regulate neurogenesis. Mol Cell 51, 349–59 (2013).

10. Vierbuchen, T. et al. Direct conversion of fibroblasts to functional neurons by defined factors. Nature 463, 1035–41 (2010).

11. Wapinski, O.L. et al. Hierarchical mechanisms for direct reprogramming of fibroblasts to neurons. Cell 155, 621–35 (2013).

12. Ang, C.E. & Wernig, M. Induced neuronal reprogramming. J Comp Neurol 522, 2877–86 (2014).

13. Voineagu, I. et al. Transcriptomic analysis of autistic brain reveals convergent molecular pathology. Nature 474, 380–4 (2011).

14. Iossifov, I. et al. The contribution of de novo coding mutations to autism spectrum disorder. Nature 515, 216–21 (2014).

15. Ronemus, M., Iossifov, I., Levy, D. & Wigler, M. The role of de novo mutations in the genetics of autism spectrum disorders. Nat Rev Genet 15, 133–41 (2014).

16. Gilman, S.R. et al. Rare de novo variants associated with autism implicate a large functional network of genes involved in formation and function of synapses. Neuron 70, 898–907 (2011).

17. Iossifov, I. et al. De novo gene disruptions in children on the autistic spectrum. Neuron 74, 285–99 (2012).

18. De Rubeis, S. et al. Synaptic, transcriptional and chromatin genes disrupted in autism. Nature 515, 209–15 (2014).

19. O’Roak, B.J. et al. Multiplex targeted sequencing identifies recurrently mutated genes in autism spectrum disorders. Science 338, 1619–22 (2012).

20. O’Roak, B.J. et al. Sporadic autism exomes reveal a highly interconnected protein network of de novo mutations. Nature 485, 246–50 (2012).

21. Hormozdiari, F., Penn, O., Borenstein, E. & Eichler, E.E. The discovery of integrated gene networks for autism and related disorders. Genome Res 25, 142– 54 (2015).

22. Meng, L. et al. Towards a therapy for Angelman syndrome by targeting a long non-coding RNA. Nature 518, 409–12 (2015).

23. Cheetham, S.W., Gruhl, F., Mattick, J.S. & Dinger, M.E. Long noncoding RNAs and the genetics of cancer. Br J Cancer 108, 2419–25 (2013).

24. Gupta, R.A. et al. Long non-coding RNA HOTAIR reprograms chromatin state to promote cancer metastasis. Nature 464, 1071–6 (2010).

25. Coe, B.P. et al. Refining analyses of copy number variation identifies specific genes associated with developmental delay. Nat Genet 46, 1063–71 (2014).

26. Cooper, G.M. et al. A copy number variation morbidity map of developmental delay. Nat Genet 43, 838–46 (2011).

27. Turner, T.N. et al. Genomic Patterns of De Novo Mutation in Simplex Autism. Cell 171, 710–722 e12 (2017).

28. Xiang, C. et al. RP58/ZNF238 directly modulates proneurogenic gene levels and is required for neuronal differentiation and brain expansion. Cell Death Differ 19, 692–702 (2012).

29. Ohtaka-Maruyama, C. et al. RP58 regulates the multipolar-bipolar transition of newborn neurons in the developing cerebral cortex. Cell Rep 3, 458–71 (2013).

30. Baubet, V. et al. Rp58 is essential for the growth and patterning of the cerebellum and for glutamatergic and GABAergic neuron development. Development 139, 1903–9 (2012).

31. Armentano, M., Filosa, A., Andolfi, G. & Studer, M. COUP-TFI is required for the formation of commissural projections in the forebrain by regulating axonal growth. Development 133, 4151–62 (2006).

32. Borello, U. et al. Sp8 and COUP-TF1 reciprocally regulate patterning and Fgf signaling in cortical progenitors. Cereb Cortex 24, 1409–21 (2014).

33. Faedo, A. et al. COUP-TFI coordinates cortical patterning, neurogenesis, and laminar fate and modulates MAPK/ERK, AKT, and beta-catenin signaling. Cereb Cortex 18, 2117–31 (2008).

34. Harrison-Uy, S.J., Siegenthaler, J.A., Faedo, A., Rubenstein, J.L. & Pleasure, S.J. CoupTFI interacts with retinoic acid signaling during cortical development. PLoS One 8, e58219 (2013).

35. Lin, F.J., Qin, J., Tang, K., Tsai, S.Y. & Tsai, M.J. Coup d’Etat: an orphan takes control. Endocr Rev 32, 404–21 (2011).

36. Job, C. & Tan, S.S. Constructing the mammalian neocortex: the role of intrinsic factors. Dev Biol 257, 221–32 (2003).

37. Tsai, S.Y. & Tsai, M.J. Chick ovalbumin upstream promoter-transcription factors (COUP-TFs): coming of age. Endocr Rev 18, 229–40 (1997).

38. O’Leary, D.D., Chou, S.J. & Sahara, S. Area patterning of the mammalian cortex. Neuron 56, 252–69 (2007).

39. Al-Kateb, H. et al. NR2F1 haploinsufficiency is associated with optic atrophy, dysmorphism and global developmental delay. Am J Med Genet A 161A, 377-81 (2013).

40. Brown, K.K. et al. NR2F1 deletion in a patient with a de novo paracentric inversion, inv(5)(q15q33.2), and syndromic deafness. Am J Med Genet A 149A, 931-8 (2009).

41. Cardoso, C. et al. Periventricular heterotopia, mental retardation, and epilepsy associated with 5q14.3-q15 deletion. Neurology 72, 784–92 (2009).

42. Malan, V. et al. Molecular characterisation of a prenatally diagnosed 5q15q21.3 deletion and review of the literature. Prenat Diagn 26, 231–8 (2006).

43. Quinn, J.J. et al. Rapid evolutionary turnover underlies conserved lncRNA- genome interactions. Genes Dev 30, 191–207 (2016).

44. Jonk, L.J. et al. Cloning and expression during development of three murine members of the COUP family of nuclear orphan receptors. Mech Dev 47, 81–97 (1994).

45. Chanda, S. et al. Generation of induced neuronal cells by the single reprogramming factor ASCL1. Stem Cell Reports 3, 282–96 (2014).

46. Treutlein, B. et al. Dissecting direct reprogramming from fibroblast to neuron using single-cell RNA-seq. Nature 534, 391–5 (2016).

47. Mall, M. et al. Myt1l safeguards neuronal identity by actively repressing many non-neuronal fates. Nature 544, 245–249 (2017).

48. Basu, S.N., Kollu, R. & Banerjee-Basu, S. AutDB: a gene reference resource for autism research. Nucleic Acids Res 37, D832–6 (2009).

49. Elling, U. et al. A reversible haploid mouse embryonic stem cell biobank resource for functional genomics. Nature 550, 114–118 (2017).

50. Zhang, Y. et al. Rapid single-step induction of functional neurons from human pluripotent stem cells. Neuron 78, 785–98 (2013).

51. Shen, Y. et al. A map of the cis-regulatory sequences in the mouse genome. Nature 488, 116–20 (2012).

52. McLean, C.Y. et al. GREAT improves functional interpretation of cis-regulatory regions. Nat Biotechnol 28, 495–501 (2010).

53. Chambers, S.M. et al. Highly efficient neural conversion of human ES and iPS cells by dual inhibition of SMAD signaling. Nat Biotechnol 27, 275–80 (2009).

54. Matsui, T. et al. Neural stem cells directly differentiated from partially reprogrammed fibroblasts rapidly acquire gliogenic competency. Stem Cells 30, 1109–19 (2012).

55. Kim, J. et al. Direct reprogramming of mouse fibroblasts to neural progenitors. Proc Natl Acad Sci U S A 108, 7838–43 (2011).

56. Qureshi, I.A., Mattick, J.S. & Mehler, M.F. Long non-coding RNAs in nervous system function and disease. Brain Res 1338, 20–35 (2010).

57. Ulitsky, I. & Bartel, D.P. lincRNAs: genomics, evolution, and mechanisms. Cell 154, 26–46 (2013).

58. Ulitsky, I., Shkumatava, A., Jan, C.H., Sive, H. & Bartel, D.P. Conserved function of lincRNAs in vertebrate embryonic development despite rapid sequence evolution. Cell 147, 1537–50 (2011).

59. Chu, C., Qu, K., Zhong, F.L., Artandi, S.E. & Chang, H.Y. Genomic maps of long noncoding RNA occupancy reveal principles of RNA-chromatin interactions. Mol Cell 44, 667–78 (2011).

60. Wang, L., Feng, Z., Wang, X., Wang, X. & Zhang, X. DEGseq: an R package for identifying differentially expressed genes from RNA-seq data. Bioinformatics 26, 136–8 (2010).

61. Leek, J.T., Johnson, W.E., Parker, H.S., Jaffe, A.E. & Storey, J.D. The sva package for removing batch effects and other unwanted variation in high-throughput experiments. Bioinformatics 28, 882–3 (2012).

62. Ernst, J. & Kellis, M. Discovery and characterization of chromatin states for systematic annotation of the human genome. Nat Biotechnol 28, 817–25 (2010).

